# Simulations of an extended Tau/tubulins interface reveal a complex disorder-disorder interplay mediated by the C-terminal tails

**DOI:** 10.64898/2026.04.30.721901

**Authors:** Jules Marien, Chantal Prévost, Sophie Sacquin-Mora

## Abstract

Building on a complex between a tubulin protofilament (PF) and a fragment of the Tau protein containing residues 169 to 367, we investigate the dynamics of the disordered elements of the system, namely the tubulin C-terminal tails (CTTs) and the Tau protein, using classical all-atom molecular dynamics simulations. Our results show that CTTs adopt a hook-like dynamic pattern on the bare PF while remaining highly mobile. The binding of Tau on the PF surface alters the dynamics of the *α*I-CTTs in a sequence-dependent manner. While the repeat domains of Tau are mostly maintained on the PF by weak and strong binding patches with the tubulin cores, the Proline-Rich Region (PRR) relies on the wrapping phenomenon of *α*I-CTTs to fuzzily stabilize its interaction with the PF. Our study thus provides a deep dive into the dynamic interplay between the Tau protein and the CTTs of microtubules, the latter being characterized extensively using a variety of disorder-adapted metrics.

**TOC Graphic:** 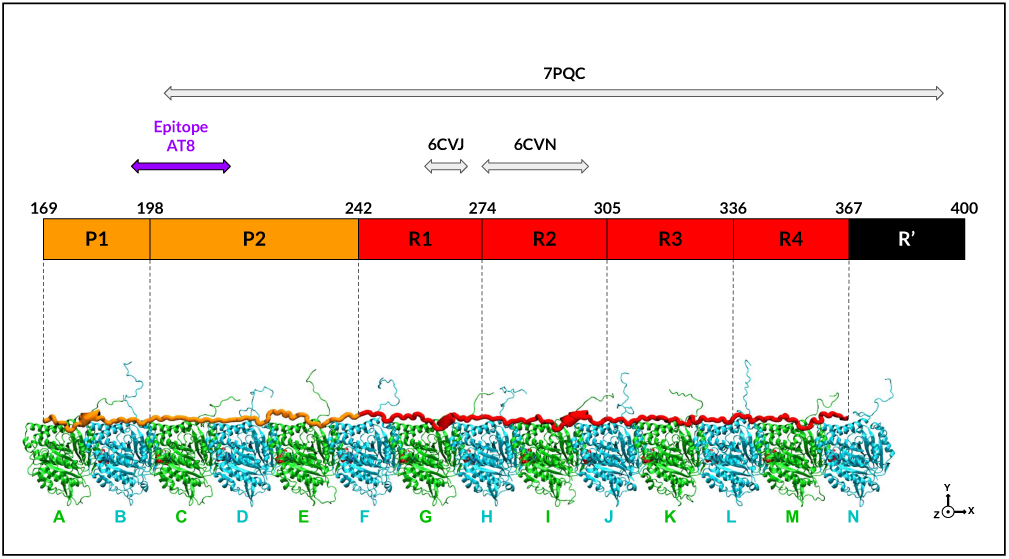

## Introduction

The cytoskeleton in neuronal axons is vital to the transport of information within the brain. In particular, microtubules (MTs) are responsible, together with actin filaments, for molecular transport along the axon.^1,2^ MTs are long, hollow, macromolecular structures, which are composed of 13 to 15 protofilaments (PFs) arranged in a helical manner. PF are themselves made of *α*-*β*-tubulin heterodimers, with various possible sequences for the *α*- and *β*- tubulins called isotypes. While some isotypes are ubiquitous in all cell types, such as *α*I tubulin and *β*I tubulin, some are cell-specific, such as the *β*III tubulin which is mostly expressed in neurons.^3^ Tubulins are structured in a globular core of around 430 amino-acids, from which a disordered C-Terminal Tail (CTT) emerges from the MT surface and interacts with the cytoplasm. Most sequence differences between tubulin isotypes are located in the CTT, which can vary in length (from 13 to 24 amino-acids) and composition. ^4^ In addition, tubulin CTTs are a target for many post-translational modifications (PTMs). The diversity induced by the various tubulin isotypes and their PTM pattern is known as forming the *tubulin code*, and is believed to encode biological functions.^4,5^

MTs growth and activity are regulated by Microtubule-Associated Proteins (MAPs). Among these, the Tau protein is the most abundant in neurons and represents over 80% of MAPs.^6,7^ Tau enhances MT growth, stabilizes MT structures and can even compact the MT lattice when Tau proteins aggregate around it to form Tau *envelopes*.^8–10^ Tau is composed of 441 amino-acids in its longest isoform and is an Intrinsically Disordered Protein (IDP) containing several regions: the projection region (residues 1 to 150), the Proline-Rich Region (PRR, residues 151 to 241), the repeat domains region (residues 242 to 400) and the C-terminal region (residues 401 to 441).^11^ Most attention has been focused on the repeat domains since they were identified to bind strongly to the MT lattice, but recent years have seen a resurgence of interest in the PRR, as it was identified to also significantly participate in interactions with tubulins.^12,13^ Furthermore, the PRR contains the AT8 epitope, a part of the Tau protein exhibiting characteristic hyperphosphorylations in Alzheimer’s disease.^14,15^ Understanding the dynamics around the AT8 epitope is thus important in order to later assess the effects of phosphorylations on the interaction between the PRR and the MT. However, little is known regarding this interface on the atomic level, due to the technical difficulties encountered by experiments when studying IDPs such as Tau, but also due to the lack of control over the tubulin isotype expression in MT, which does not allow to obtain a clear picture of the dynamic interplay at the scale of a single molecule.

In 2021, Brotzakis et al. published an all-atom simulation study on the interaction between a PF and a Tau fragment ranging from residues 202 to 395 (almost half of the PRR and all repeat domains).^16^ Their work revealed the presence of alternating weak and strong binding patches within the repeat domains interacting with the tubulin cores. Modelled residues from the PRR were found to interact only transiently with the PF surface, but a part of the AT8 epitope was truncated.

In the present work, we start from the equilibrated complex deposited in the Protein Data Bank^17^ (PDB) by Brotzakis et al., that we extend to include the P1 domain of Tau (see Figure 1). We perform 1*µ*s-long all-atom classical molecular dynamics simulations of *α*I/*β*I and *α*I/*β*III PFs bare or in complex with a fragment of Tau encompassing residues 169 to 367. From the bare PFs, we extract the dynamic profile of each CTT isotype, and show how it is disturbed upon addition of Tau. We observe that the interaction of the *α*-CTTs with Tau is region-dependent and that the PRR mostly relies on the previously described ‘wrapping phenomenon’ of the CTTs around Tau for stabilization of the interface.^18,19^ Finally, the comparison between bare PFs and bound PFs also allows to observe the compaction of the MT lattice by Tau.

**Figure 1:**
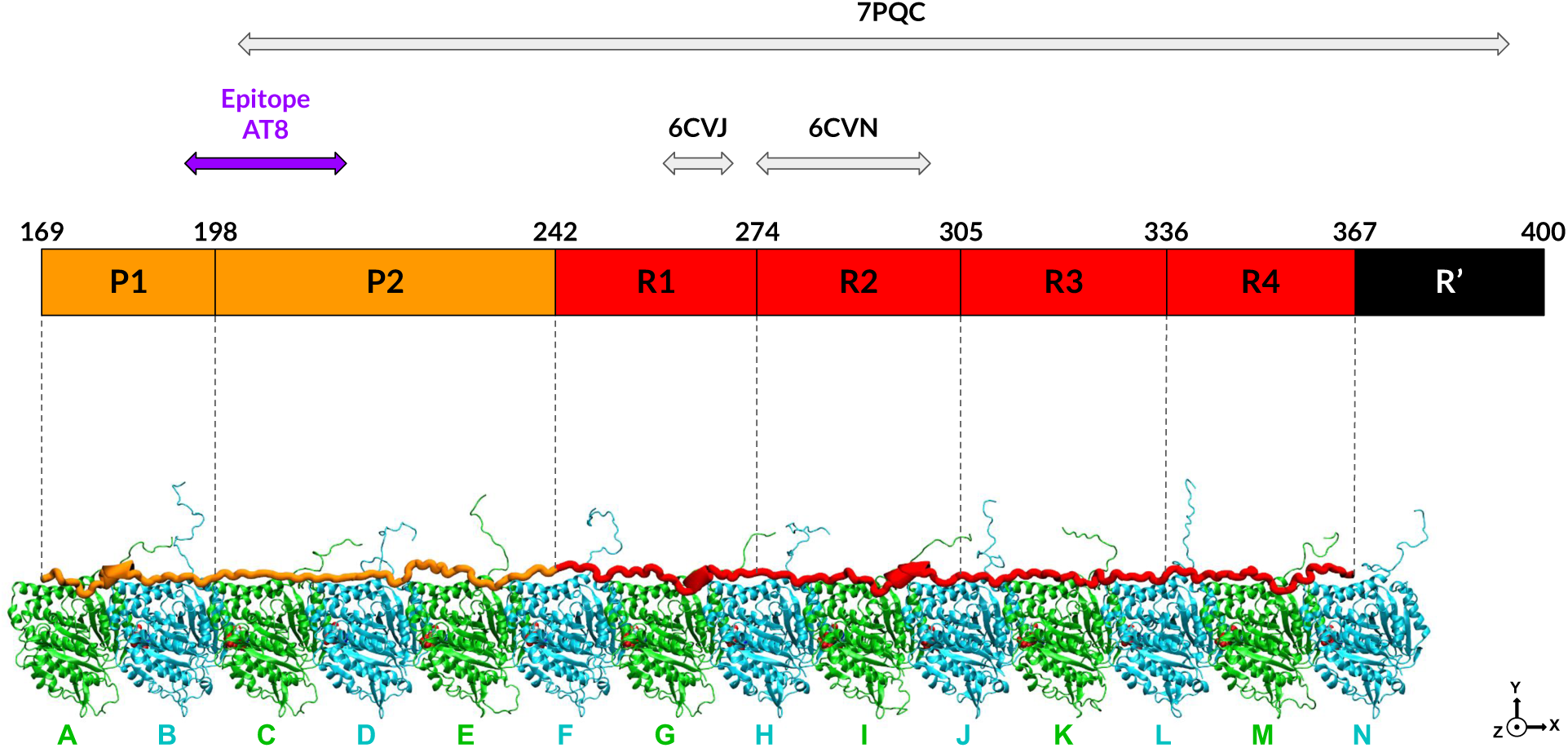
Representation of the *α*I/*β*III PF in complex with the Tau fragment spanning residues 169 to 367. *α*-tubulins are in green, *β*-tubulins in blue. GTP, GDP and Mg^2+^ cations are represented at the intertubulin interfaces in Van der Waals representation. The modelled part of the Proline-Rich Region (PRR) is in orange, the R1/R2/R3/R4 repeat domains are in red. Tubulin cores and their C-terminal tails (CTTs) are in NewCartoon representation. The molecular structure was rendered using VMD (Visual Molecular Dynamics).^20^

## Material and Methods

### 1. Building the complex

We started from the work by Brotzakis et al. and the deposited PDB structures 7PQC and 7PQP to build our systems.^16^ These structures contain seven *α*/*β* dimers in complex with a Tau fragment spanning residues Ser202 to Lys395. However, we wanted to probe the dynamics of the PRR, and notably of the AT8 epitope of Tau (residues 194 to 211), so we had to extend the system towards the N-ter extremity of Tau.

In the 7PQC model, the Tau chain is tightly bound to the tubulins from Lys240 to Gly367 and is described by the authors as representative of a *ground state* of binding. However, the rest of the N-ter part (Ser202 to Lys240) only weakly binds the surface of the PF, which would make its extension difficult. We thus considered the 7PQP model for the Ser202 to Lys240 part. The P2 domain of Tau is bound to the tubulin cores in this structure, so we decided to combine both 7PQC and 7PQP models in a hybrid starting structure. To do this, we placed both disjointed parts of Tau in a single PDB file, then added the tubulins of the 7PQC model and performed a minimization step with NAMDv2.13 to close the gap.^21^ All atoms but those of the 237-SSAKSR-242 part were harmonically restrained, then 10000 steps of minimization were performed using the CHARMM36m forcefield. ^22^ The output was a hybrid model where all modelled residues of Tau are close to the PF surface and aligned with its longitudinal direction, which we defined as the X direction.

We then used the Heligeom tool^23^ in order to extend the PF in the N-ter direction of Tau. We created a structure containing two tubulin dimers made of the first four monomers of the hybrid model, which we then provided to the Heligeom website^24^ by labelling the first tubulin dimer as a single chain and the second dimer as another. Heligeom was initially designed to study and extend helicoidal geometries of binding, but it turns out that it can also extend longitudinal periodic geometries since they are an helicoidal geometry with an infinite pitch. We required Heligeom to extend the PF structure by two dimers. CTTs were ignored when generating the helical transformation. The resulting extended PF was fused with the hybrid model so as to add only three tubulin monomers in the N-ter direction of Tau.

The next step was to extend the Tau fragment to Thr169 on the three additional tubulin monomers, for which we used the Modeller tool.^25^ We provided as template the fragment of Tau ranging from Asp283 to Leu315 and the four monomers of tubulin it is binding, and the Arg211 to Leu215 Tau fragment with the four monomers of tubulins containing the ones added by Heligeom. These templates ensured that the generated extension is aligned with the already existing structure of Tau and that the added portion remained close to the tubulin cores. One hundred models were generated differing only by the structure of the Tau fragment. We selected the one with the best DOPE score (which is the internal score of MODELLER). This extension of Tau was then added to the system.

The resulting structure constituted an overall template of the system, but we still needed to modify the tubulin sequences used by Brotzakis et al. for the ones we used in our previous studies and add the corresponding CTTs.^18,19^ We used Modeller again to do so, using sequences from sheep for the *α*I tubulin (gene TUBAIA, Uniprot accession number D0VWZ0) and for the *β*III tubulin (gene TUBB3, Uniprot accession number Q13509). One hundred models were generated, and we selected the one with the largest cumulative radii of gyration of the CTTs in order to facilitate equilibration, as we had observed in previous simulations that CTTs tend to extend at first when the equilibration is performed.^18,19^ Another system was built with *β*I tubulin (gene TUBB, Uniprot accession number W5PPT6) instead of *β*III for the PF, using the *β*III model as a template to get as close a starting conformation as possible for the two types of PF (*α*I/*β*I and *α*I/*β*III).

As a result, the obtained complexes presented a PF of seventeen tubulin monomers. We decided to delete the *α*/*β*/*α* tubulin monomers located on the C-terminal side of the Tau fragment in order to reduce computational costs and maintain dimer parity. Since the pseudorepeat of Tau (residues Asn368 to Lys395) was then no longer in contact with tubulins, we truncated it from the system. Associated GTP/GDP nucleotides and Mg^2+^ cation were also removed. The final Tau fragment studied here spans residues Thr169 to Glu367, which correspond to most of the P1 domain, the P2 domain and the R1/R2/R3/R4 domains, and has a starting conformation close to the tubulin cores along all of its sequence. A representation of the *α*I/*β*III PF in complex with Tau is shown on Figure 1. In addition to simulations of the PF in complex with the Tau fragment, we also removed the Tau fragment to obtain seven-dimer-long bare *α*I/*β*I and *α*I/*β*III PFs in order to study the dynamics of free CTTs. The detailed composition of the four systems is available in Table SI-1.

### 2. Molecular dynamics simulations

All four systems (bare *α*I/*β*I PF, *α*I/βI PF in complex with the Tau fragment, bare *α*I/*β*III PF, *α*I/*β*III PF in complex with the Tau fragment) were solvated in a periodic box of dimensions 670 Åx140 Åx140 Å using CHARMM-GUI,^26^ where the longer direction X corresponds to the longitudinal axis of the PF. Systems were neutralized using KCl ions and a concentration of 0.15 mol.L*^−^*^1^ was added. Simulations and restraints were carried out using the NAMDv2.14 software.^21^ The CHARMM36m forcefield was used to describe the complex (proteins, GTP, GDP, Mg^2+^) with its associated TIP3P water model.^22^ Ionic parameters are standard CHARMM ions from Beglov and Roux.^27^ The Particle Mesh Ewald scheme was employed for electrostatic calculations with a 12 Å cutoff and a 10 Å switching, and an interpolation order of 6. ^28^ The temperature was set to 300 K through Langevin dynamics with a damping of 1 ps*^−^*^1^. The integration timestep was set to 2 fs and the SHAKE algorithm was used to constrain covalent hydrogen bonds.^29^

A 10 000-step minimization was performed followed by a 200 ps equilibration in the NPT ensemble in order to effectively pressurize the system at 1 atm. All heavy atoms besides those of the solvent were restrained by a harmonic potential with a force constant of 1 kcal.mol*^−^*^1^.Å*^−^*^2^. The Langevin piston scheme was employed as a barostat with a 1000 fs period and 500 fs decay in order to accommodate for the size of the system. This was followed by a 1 ns equilibration in the NVT ensemble to allow the solvent to properly equilibrate. The rest of the equilibration was performed in the NPT ensemble with a Langevin piston of 100 fs period and 50 fs decay. Then, restraints were gradually lowered on the heavy atoms to let the complex relax. 500 ps were spent with full restraints, then 1 ns with ten times weaker restraints, then another 1 ns with a hundred times weaker restraints.

Then, we implemented harmonic constraints on the Y and Z directions (perpendicular to the axis of the PF) in order to keep the PF from bending or drifting in the box. We constrained the *C_α_* of Phe169 in all *α*-tubulin monomers and the equivalent Phe167 in all *β*-tubulin monomers as this residue is located close to the center of mass of the tubulin core. Two successive 1 ns steps were performed with no other restraints, resetting the center of the restraints between each step to allow the complex to accommodate for the constraints and to relax disordered regions such as the Tau fragment and the CTTs. This careful equilibration scheme also allows for tubulins to relax from the constrained conformations they had adopted from the simulation protocol they originated from, as in the work of Brotzakis et al. tubulin cores were simulated with restraints imposed by a cryo-EM map of a microtubule.^16^

For the production step, Y and Z constraints were extended to *C_α_*s of residues 168-169-170 for *α*-tubulin and the equivalent residues 166-167-168 for *β*-tubulin. Each complex was simulated for 1000 ns, sampling frames every 100 ps.

### 3. Analyses

Analyses of the trajectories were performed using the MDAnalysis and MDTraj python libraries.^30,31^ The Proteic Menger Curvatures (PMCs), Local Curvatures (LCs) and Local Flexibilities (LFs) metrics were computed using the Menger_Curvature MDAKit.^32^ In this paper, the Root-Mean-Square Fluctuation (RMSF) is defined as the fluctuation of the Tau fragment with regards to its conformation in the first frame. The alignment of the structure is performed on the tubulin cores, not on the Tau fragment itself, so this modified RMSF also reports on potential displacements of Tau with regards to the original interface. Contact between two residues is defined as any of their heavy atoms being within 5 Å of each other. Plots were displayed using the Matplotlib python package,^33^ and structure illustrations were produced with the Visual Molecular Dynamics (VMD) software.^20^ Unless specifically specified, we only considered tubulin monomers C to L in the analyses. Since dimers AB and MN are at the extremities of the complex and the Tau fragment does not fully extend onto them, removing them from analyses should help avoid truncation effects arising from the incompleteness of the Tau fragment.

## Results and Discussion

### 1. In-depth characterization of the C-terminal tails (CTTs) on a protofilament (PF)

#### a. Free CTTs

We first characterize the dynamics of the CTTs without the Tau fragment in order to obtain a profile of how a *free* CTT behaves, and whether there is a difference between the isotype combinations (*α*I/*β*I vs *α*I/*β*III). For this purpose, we considered the two simulations without the Tau fragment. For *α*-CTTs, we studied the behavior of monomers C, E, G, I, and K, and for *β*-CTTs of monomers D, F, H, J and L. As a first approach, we made the assumption that all CTTs are equivalent in these systems since their direct monomeric environment is the same. We thus concatenated their contact pattern into a single map per type of CTT, essentially treating each CTT as if 5 replicas of 1*µ*s had been performed (Figure 2). *α*I-CTTs mostly display contacts at their base with residues from the tubulin cores (residues *α*-Asp345, *β*-Arg391, Figure 2). A difference exists between *α*I-CTTs in complex with *β*I tubulins and *α*I-CTTs in complex with *β*III tubulins. Residue *α*-Arg339 is located on a loop facing the *α*-helix preceding the CTT of the tubulin, and for *α*I-CTTs in complex with *β*III tubulins, contacts Glu447 less than 10% of the time. For *α*I-CTTs in complex with *β*I tubulins, Glu445, 446 and 447 all make contact with *α*-Arg339 for more than 20% of the time. These differences may imply that the dynamics of the *α*I-CTT is sensitive to the isotype of its neighboring monomer. If this is the case, the contact map indicates that this is not due to strong interactions between *α*- and *β*-CTTs since no residues from the *α*-CTT appear in the *β*-CTT map and vice versa. As for the *β*I- and *β*III-CTTs, they both contact Arg306, Arg309, Lys336, Ser339, and Trp344 on the *β*-tubulin core and Lys401 on the *α*-tubulin core. This contact pattern is in full agreement with previous simulations of tubulin dimers by Laurin et al.^34^ This similar behavior despite the use of a different forcefield reinforces our confidence in the sampling and the dynamics of the disordered parts of the complexes.

**Figure 2:**
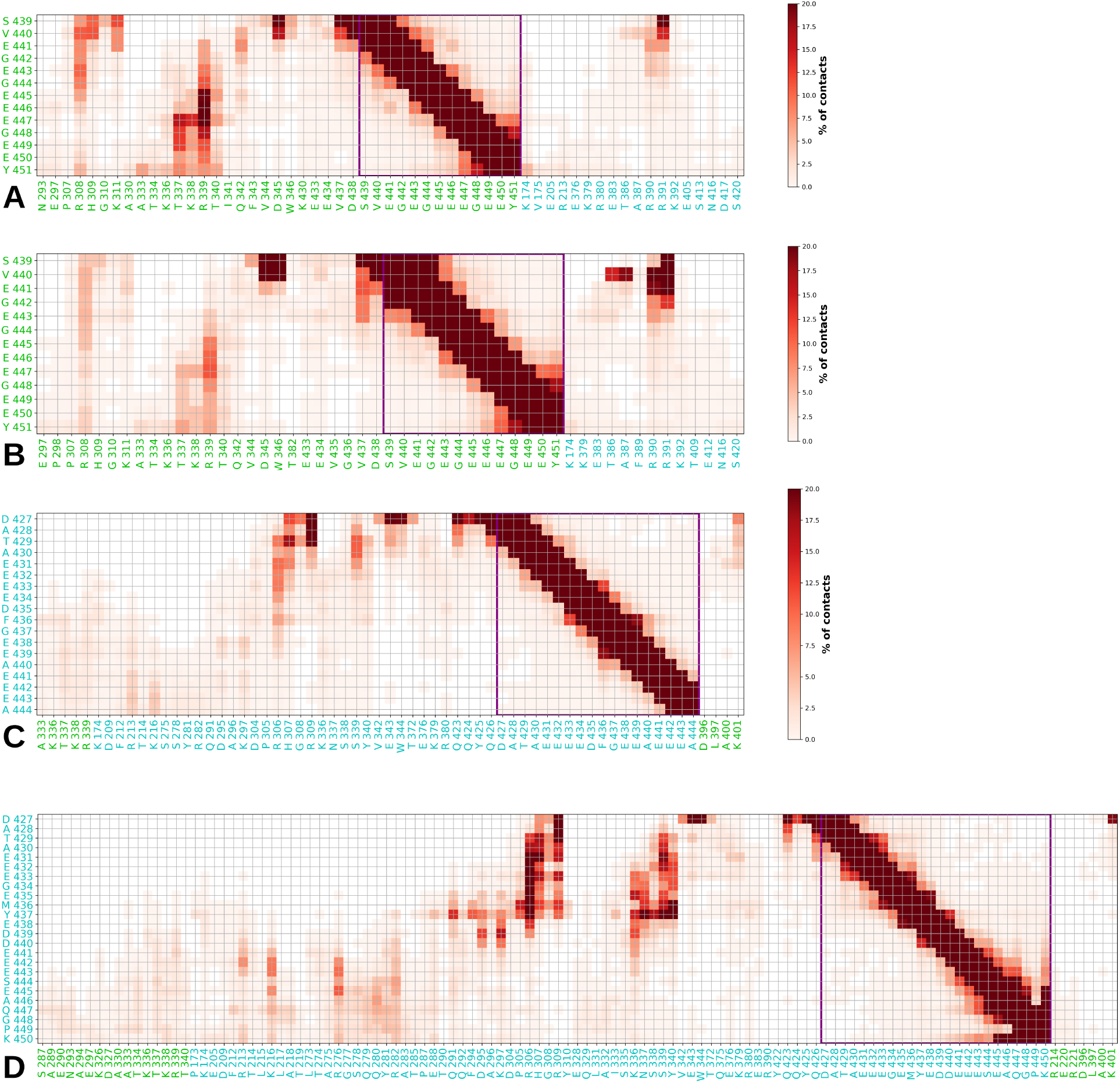
Concatenated contact map between the CTTs and tubulin *α* (in green) and *β* (in blue) in absence of the Tau protein. Tubulin residues that did not contact the CTT at least once during the simulations were filtered out. A) *α*I CTTs in the *α*I/*β*I complex. B) *α*I CTTs in the *α*I/*β*III complex. C) *β*I CTTs. D) *β*III CTTs

The intramonomer interactions of the residues of a CTT with itself are clearly indicative of the fuzziness of CTTs. No long-lasting contact between residues farther than six residues appart was formed within the CTTs. Interestingly, for *α*I and *β*III CTTs, longer-lasting contacts are observed for the six terminal residues. This indicates a certain propensity of these residues to adopt a hook-like conformation. This can be confirmed and further precised by looking at the LC profiles of CTTs (Figure 3). For *α*I-CTTs, the curvature slightly increases for the 2 last residues Gly448 and Glu449 compared to the previous residues. For *β*III CTTs, the curvature strongly increases for residues Ala446, Gln447 and Gly448, in agreement with the strong hook-like contacts spotted on the contact map (Figure 2D).

**Figure 3:**
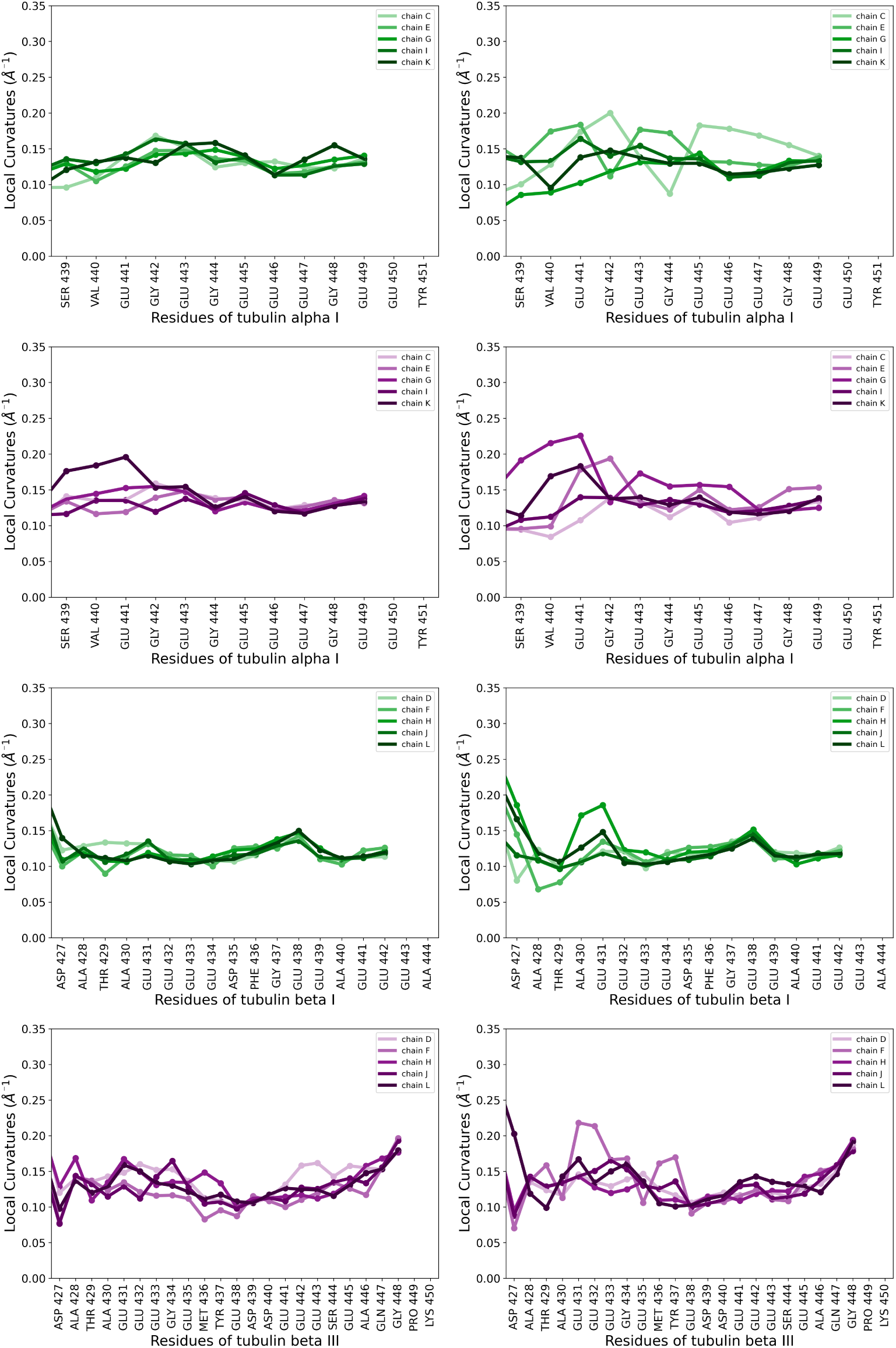
Local Curvatures (LCs) of the CTTs with (right column) and without (left column) Tau. First row: *α*I CTTs in the *α*I/*β*I complex. Second row: *α*I CTTs in the *α*I/*β*III complex. Third row: *β*I CTTs. Fourth row: *β*III CTTs

LCs and LFs also reveal interesting features of the dynamics of free CTTs (Figures 3 and 4). For reference, an *α*-helix displays LC values of 0.30 Å*^−^*^1^ while residues in a *β*-sheet are usually below 0.05 Å*^−^*^1^ (see Ref^32^), and the central residue of a very flexible polyalanine peptide has a LF value of 0.07 Å*^−^*^1^.^35^ Most LC values for the CTTs observed here are comprised between 0.10 and 0.15 Å*^−^*^1^, and most LFs values are comprised between 0.04 and 0.08 Å*^−^*^1^, which is indicative of a fully disordered behavior.^35^ The curvature profiles of *α*I-CTTs show a slight decrease at Glu446 for the *α*I/*β*I system, and at Gly444 when in the *α*I/*β*III system. For *β*I-CTTs, a peak is universally observed at Glu438, which would translate as a tendency to mark a turn at this residue, a conformation propensity that would resemble a loose hook. On the other hand, *β*III-CTTs have the least converged behavior, probably due to their longer sequence. All five CTTs still reach an agreement and a lower LC value at Asp439. The sharp increase in LC values for the terminal residues would translate in a conformational propensity for a tight hook.

**Figure 4:**
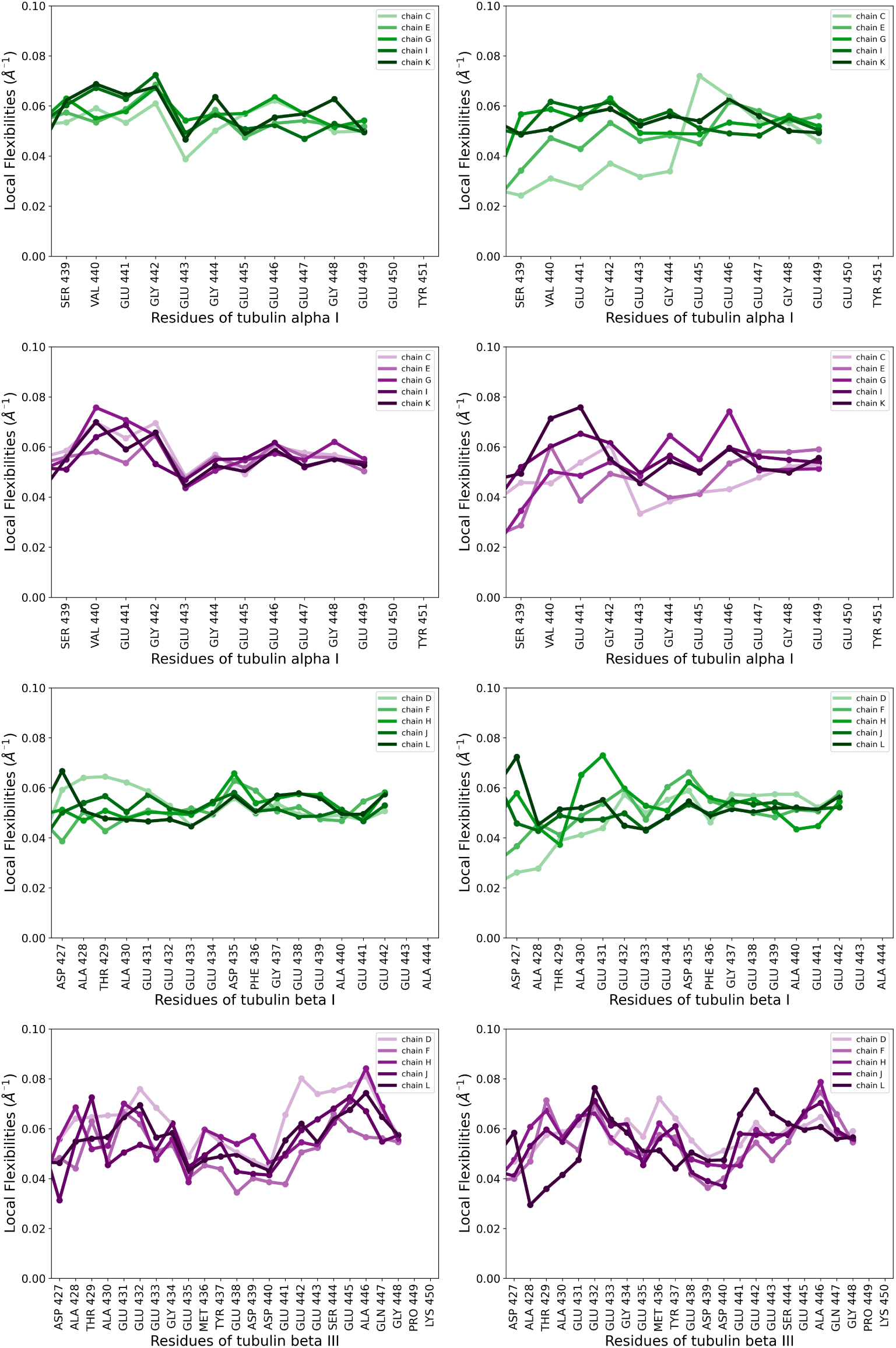
Local Flexibilities (LFs) of the CTTs with (right column) and without (left column) Tau. First row: *α*I CTTs in the *α*I/*β*I complex. Second row: *α*I CTTs in the *α*I/*β*III complex. Third row: *β*I CTTs. Fourth row: *β*III CTTs

LF profiles provide information on the flexibility experienced by the chain around a given residue. For the *α*I CTTs, LFs values are particularly high at their base then plummet at Glu443. This residue does not interact significantly with others according to the contact maps (Figures 2A-B), which would indicate that the lower LF is due to longer range effects of the CTT dynamics. *β*I-CTTs experience a relatively flat profile of LFs with a slightly higher flexibility at Asp435. Similarly to the LCs, the LFs of *β*III-CTTs are not as converged as for the other types of CTTs, but trends can still be read in their profile. A common drop in flexibility resembling the one of *α*I is found at Glu435, followed by a lower flexibility region up to Asp440, after which residues experience a high flexibility up to Ala446, which becomes lower for Gln447 and Gly448. These LF profiles are in agreement with a hook-like behavior of the *β*III termini, with Ala446 probably acting as the turning point.

CTTs are often referred to as *E-hooks* for their high content in glutamate residues and their ability to recruit other MAPs, notably dynein and kinesin. ^36–39^ According to our results, this name might actually be fitting in a much more literal sense. Data contained in Figures 2, 3 and 4 indeed hints at different hook properties between the free CTTs of isotypes *β*I and *β*III: a loose hook for the former, and a tight hook for the latter. The disordered nature of the CTTs captured in these Figures is also an experimental reality as was directly measured with NMR by Wall et al.,^40^ and as was computed by the DSSP algorithm in Figure SI-1.

LCs and LFs profiles of the free CTTs also show that the dynamics of the five CTTs of each category has mostly converged. For the LCs (Figure 3), there is indeed little difference between each CTT except a slight curvature increase at the base of CTT-*α*I of monomer K and at the Glu442-Glu443 for CTT-*β*III of monomer D. Likewise, the LFs profiles are very similar between chains for *α*I- and *β*I-CTTs, and still very comparable for *β*III chains, which show more variability probably due to their length. This implies that while the simulations cannot be considered as fully converged, the sampling is still extensive enough to significantly probe CTT dynamics. It also means that differences that will be spotted in presence of the Tau monomer can be considered meaningful.

#### b. Impact of Tau binding on the CTT dynamics

Next, we considered the two simulations (*α*I/*β*I PF and *α*I/*β*III PF) where the Tau fragment is bound on the PF. We investigated the changes induced in CTT dynamics by reproducing the analyses discussed in the previous part. The difference between the concatenated contact map for free CTTs and for CTTs in the presence of Tau is represented on Figure SI-2. For the *α*I-CTT, the presence of Tau induces significant contact losses (more than 10%) for residues Glu446/Glu447 with residue *α*I-Arg339, and contact reinforcements for the base of the CTT with residues Arg390/Arg391 of both *β* isotypes. Intra-CTT contacts remain similar for the *α*I-CTT in the *α*I/*β*III system, and only a 5% increase is spotted for residues Glu443/Gly444 with Glu447 for the *α*I-CTT in the *α*I/*β*I system. *β*I-CTTs keep their self-interaction contacts, but experience increased contact time with *β*I-Trp344 and residues *β*I-Arg306 to Arg309. *β*III-CTTs loose nearly all the contact time that their base had with the *β*III tubulin core, while residues *β*III-Thr429 to Met436 increase their contact time by around 10%. The presence of Tau therefore has an impact on the interaction of tubulin CTTs with the core and on the interactions of *β*III-CTTs with themselves.

Another intriguing effect of the presence of Tau is the appearance of a helical structure for 10% of the time for residue *β*-Asp427 for both *β*I and *β*III isotypes (see Figure SI-1). This is confirmed by the increase in contact time observed between *β*-Asp427 and *β*-residues 423-QQY-425 in both cases (see Figure SI-2). *β*-Asp427 is located at the very base of the *β*-CTT, at the junction with the last *α*-helix of the tubulin core. The stabilization and extension of this helix by the presence of Tau is interesting since it could be related to the stabilizing MAP function of the Tau protein for the microtubule.

We next focused on the impact on the local dynamics of the CTTs by computing their LCs and LFs profiles in presence of Tau (Figures 3 and 4). For all CTT isotypes, disruptions are observed at the base residues of CTTs for certain monomers. For the *α*I-CTTs, monomers C and E for the *α*I/*β*I system and monomers C, E and G are impacted by the presence of Tau, which translates into an increase of the curvature at the base and a decrease in flexibility. For the *β*I CTTs, the same is observed for monomers D and F. For the *β*III CTTs, only monomer F displays higher curvatures at Glu442 and Tyr437, while the base of monomer L has a smaller flexibility. Based on our previous simulations of the CTTs of tubulin in contact with a Tau fragment, we immediately investigated whether these differences could be caused by the already characterized *wrapping phenomenon* of the CTTs around Tau described in our previous works.^18,19^

#### c. CTTs and Tau : it’s an (uneven) wrap !

We conducted a visual inspection of both simulations for all monomers C to L in order to identify whether *wrapping* events had been sampled. Events of interest were recorded in Tables SI-2 to SI-5. A first surprising observation was that almost no *wrapping* was observed for *β*-CTTs, in contradiction with what had been found in our earlier studies. ^18,19^ For these larger PF complexes, both *β*I- and *β*III-CTTs tend to remain extended in the bulk or to interact transiently with the side of the tubulin core opposite to Tau. The second observation was that *α*-CTTs do not only wrap around the Tau monomer but also alongside it (see Figure 5 for an example with the CTT of monomer E), probably due to their short length (13 residues). This *longitudinal wrapping* can transition into a *transversal wrapping*, and vice versa, it could thus be considered as both aspects of the same interaction type. We tried to categorize both behaviors as best as possible but this dynamic phenomenon is more of a continuum than a clear-cut divide, due to how transient and dynamic the interactions of the CTTs with Tau are. The last surprising observation was that *α*-CTTs of monomers C and E wrap the Tau fragment significantly more than those of the other monomers. Since no difference was observed in the dynamics of the CTTs when the complex does not involve Tau, this difference was unexpected and worth investigating further. Out of the 20 000 frames collected for each monomer on the *α*I/*β*I and the *α*I/*β*III systems, we observed that the *α*-CTT of monomer C is wrapped transversally and/or longitudinally on the Tau fragment for 16 180 frames, and the *α*-CTT of monomer E for 16 670 frames. Other *α*-CTTs wrap barely a quarter of the time, at most.

**Figure 5:**
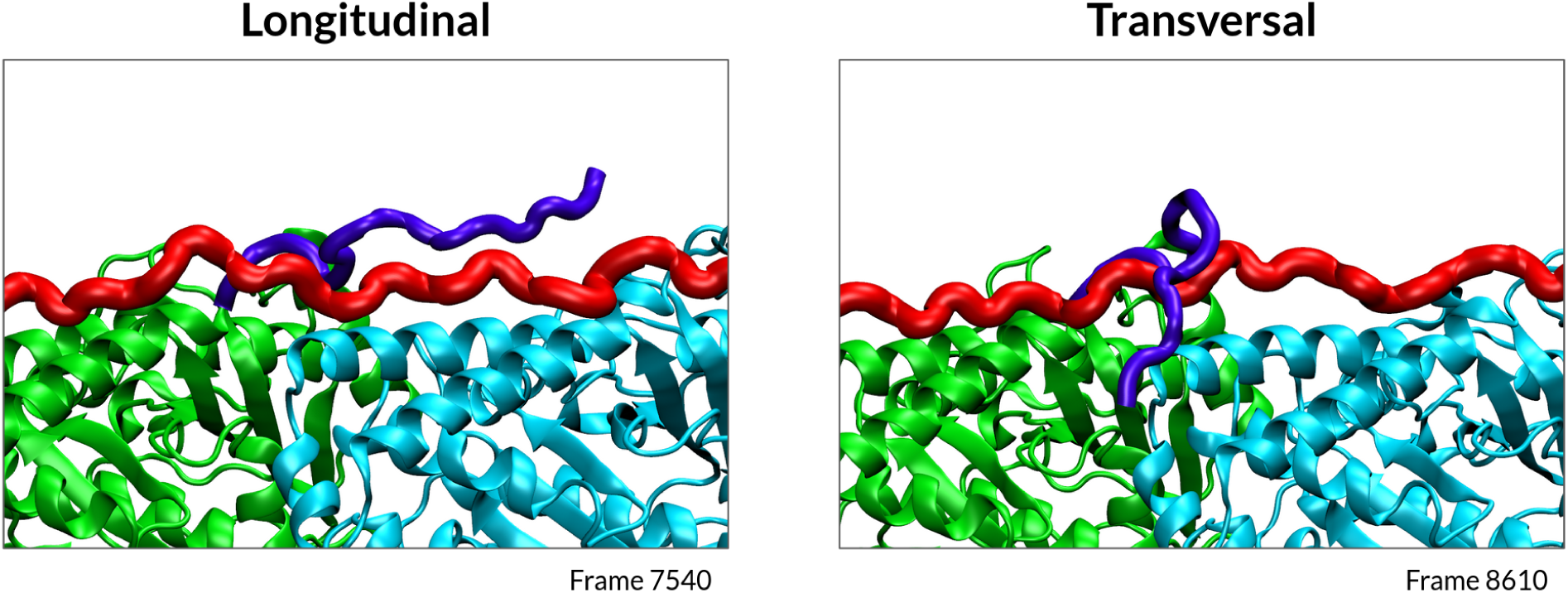
Longitudinal (left) vs transversal (right) *wrapping* of the *α*I-CTT (in purple) of monomer E on the Tau protein (in red).

We represented the PMCs of the CTTs of monomers C and E as a function of time for the second half of the trajectories, and compared their profiles between free CTT and Tau-bound CTT (Figures SI-3 and SI-4). The impact of the wrapping on the dynamics of the CTTs is clearly visible. Without Tau, *α*-CTTs C and E experience transient variations of their curvatures homogeneously along their sequence, while with Tau the PMC of each residue tends to remain fixed. For the *α*-CTT of monomer C in the *α*I/*β*I system, Gly442 and Glu445/446 remain highly curved while Gly444 is very extended, separating the CTT in two distinct sections. A similar profile can be observed for the *α*-CTT of monomer C in the *α*I/*β*III system. For the *α*-CTT of monomer E in the *α*I/*β*I system, it is Gly442 which remains extended while Val440/Glu441 and Glu443/Gly444 are very curved. In the *α*I/*β*III system, the *α*-CTT of monomer E is very curved at Glu441 and Gly442 while the rest of the residues experience different curvatures over time, although they evolve slower than for the equivalent free CTT. Overall, the *wrapping* phenomenon induced by the presence of Tau tends to increase and stabilize the curvature profile of the residues at the base of the CTTs. A noticeable finding is the specificity of the PMC profile of the glycine residues upon Tau addition. One can indeed wonder whether they might act as a dynamic flexible joint for the CTT sequence. An interesting experiment to perform based on these results would be the mutation of the glycine residues to bulkier residues among the aliphatic amino acids (valine, isoleucine, leucine) to see whether the CTTs still manage to wrap, and how the mutation would affect the binding of Tau to the microtubule.

Whether the *wrapping* phenomenon can be or even has been already observed experimentally is hard to assess, since direct-observation techniques such as microscopy or cryo-EM are not yet able to resolve the fuzziness of CTTs. However, one could hypothesize that the dynamic binding of the *α*-CTTs on Tau may look like the one experienced by MAP7 with tubulin CTTs according to solid-state NMR experiments performed by Luo et al.,^41^ although MAP7 takes an *α*-helical structure on the studied sequence^42^ and likely gets bound on the millisecond timescale.^41^

The clear divide in wrapping events between *α*-CTTs of monomers C/E and monomers G/I/K must arise from a Tau-specific interaction since these CTTs all behave similarly when free of Tau. We remark on Figure 1 that monomers C and E are bound to the PRR region of Tau while monomers G, I and K interact with the repeat domains. We therefore studied the dynamics of the Tau fragment on the PF in order to assess whether the *wrapping* discrepancy of the *α*I-CTTs could be explained by an interaction specificity for the different Tau regions.

#### d. Finding collective variables that capture the wrapping phenomenon

The impact of Tau on the dynamics of the CTTs was explored in our previous work by projecting the center of mass (COM) of the CTTs in a spherical plane (see Figure SI-8 of Ref^18^). This representation allowed to observe differences between the behavior of free CTTs and CTTs within the vicinity of the R2 repeat domain of Tau, but was not precise enough to allow for a clear characterization of the wrapping phenomenon. We thus set to find collective variables (CVs) that would reflect the longitudinal to transversal landscape of the *wrapping* identified by our observations.

The first variable Ψ*_CTT_* that we propose allows to assess whether a wrapping takes place (Figure 6, left). Projecting on the YZ plane, Ψ*_CTT_* is defined as the angle between the vector “base of the CTT/COM of the tubulin core” and the vector “base of the CTT/tip of the CTT”. This angle can be compared to the range of values Ψ*_Tau_* taken by the angle between the vector “base of the CTT/COM of the tubulin core” and the vector “base of the CTT/nearest *C_α_* of Tau”. If Ψ*_CTT_* is of the order of Ψ*_Tau_*, meaning between 220° and 300°, one can consider that the CTT is effectively wrapping around the Tau fragment. The second variable, *D_b_*_2*t*_, is designed to capture whether the wrapping is rather longitudinal or transversal (Figure 6, right). By calculating the projection of the vector “base of the CTT/tip of the CTT” onto the X axis, one can assess how elongated the CTT is along the Tau fragment.

**Figure 6:**
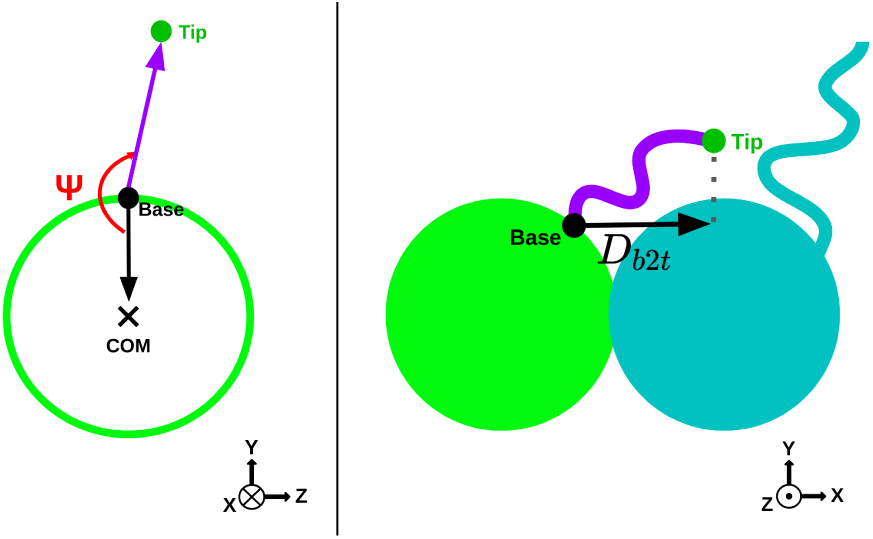
Schematic representation of the collective variables (CVs) defined in order to characterize the dynamics of the CTTs, in particular their *wrapping* propensity. Ψ*_CTT_*is represented on the left, and *D_b_*_2*t*_ on the right. Tubulin cores are green or blue circle, the CTT is in purple and the base of the CTT is represented by a black dot and its tip by a green dot.

Considering the Ψ*_CTT_* /*D_b_*_2*t*_ plane yields a characterization for each CTT of how it explores the space above the PF (Figure 7). *α*I-CTTs on the bare PF tend to either explore what would be the exposed surface of the PF in a microtubule (Figure 7a around 180°, monomers E, I and K) or the part buried at the inter-PF interface (Figure 7a between 0° and 100°, monomers C and G). The presence of Tau drastically modifies the *α*I-CTT dynamics, highlighting the high frequency of the wrapping phenomenon as the CTTs mostly explore values of Ψ*_CTT_* close to those of Ψ*_Tau_*. The several basins at different values of *D_b_*_2*t*_ tend to suggest that the CTTs transition between longitudinal and transversal wrapping modes, as was noticed by visual inspection. *β*III-CTTs on the other hand behave very similarly by only exploring the buried interface of the bare PF (Figure 7b between 0° and 100°). Addition of the Tau fragment does not modify the dynamics except for monomer L, indicating that no wrapping is taking place.

**Figure 7:**
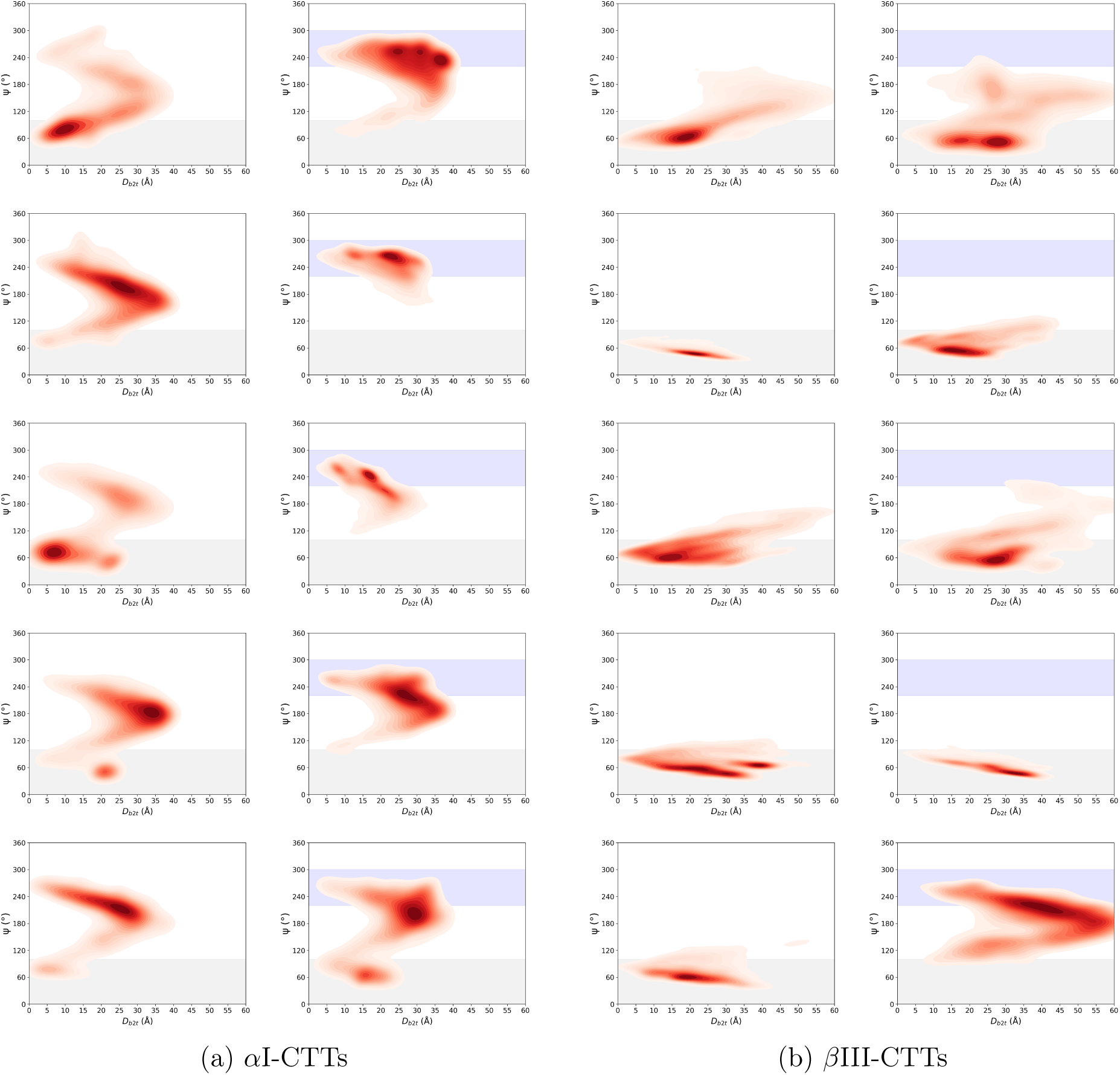
Landscapes of the Ψ*_CTT_* /*D_b_*_2*t*_ values for the CTTs of the bare *α*I/*β*III PF (left columns of 7a and 7b) and the PF in complex with the Tau fragment (right columns of 7a and 7b). From top to bottom, plots correspond to monomers C, E, G, I and K for the *α*I-CTTs and monomers D, F, H, J and L for the *β*III-CTTs. The range of Ψ*_Tau_* values is shaded in blue. The range of Ψ*_CTT_* values corresponding to a position that would be buried into the inter-PF interface is shaded in grey.

Comparable results are achieved for *α*I-CTTs of the *α*I/*β*I PFs (see Figure SI-5) as for those of the *α*I/*β*III PFs. The *β*I-CTTs however do not behave like the *β*III-CTTs. *β*I-CTTs in the bare PF can adopt values of Ψ*_CTT_* around 180° which are barely explored by the *β*IIICTTs. This difference in behavior can be explained by the shorter length of the *β*I-CTTs, but also by the absence of a lysine at their tip which is characteristic of the *β*III-CTTs and must affect the exploration of the PF surface. The presence of Tau does not strongly alter the exploration landscapes of the *β*I-CTTs, and does not cause them to engage in *wrapping*. Overall, the use of the collective variables Ψ*_CTT_* and *D_b_*_2*t*_ allows for a useful characterization of the dynamics of each individual CTT, revealing whether they engage in wrapping with Tau and which part of the PF surface is mostly explored. However one can question the ability of the Ψ*_CTT_* /*D_b_*_2*t*_ landscapes to provide a clear-cut divide between longitudinal and transversal wrappings, as it seems that these states should be interpreted more as both extremes of a continuum of binding poses rather than a binary positioning of the CTT over Tau. Nevertheless, the Ψ*_CTT_* /*D_b_*_2*t*_ landscapes provide us with uselful information when investigating the dynamics of simulated CTTs, even though further works should be carried out in order to deepen our understanding of their motion.

### 2. Tau/tubulins interface

#### a. The contact pattern is region-specific

We first studied the interface between the tubulins and Tau using a contact map representation (Figures 8 and SI-6). Tubulin monomers A, B, M and N were also included in this analysis in order to sample the entirety of the contacts made by the Tau fragment with the PF. The clear patterning of the interactions across the tubulin dimers reflects the overall extension of Tau on the PF along the trajectories. In their simulations of a similar complex, Brotzakis et al. identified strong binding patches in the repeat domains around their SKXGS motifs and weak binding patches around the N-terminal parts of the repeats.^16^ We observe the same behavior here, with the SKXGS motifs of repeats R2, R3 and R4 binding nearly at all times to their respective *α*-tubulin core and displaying an additional strong interaction pattern with *α*-tubulin residues Tyr262 to Asp345. The strong binding patches of these three repeats are also strongly bound to *β*-tubulin residues Arg390/Arg391/Lys392, three residues located at the *α*/*β* interface (residues below the green Van der Waals sphere of the *α*-CTT on Figure 9).

**Figure 8:**
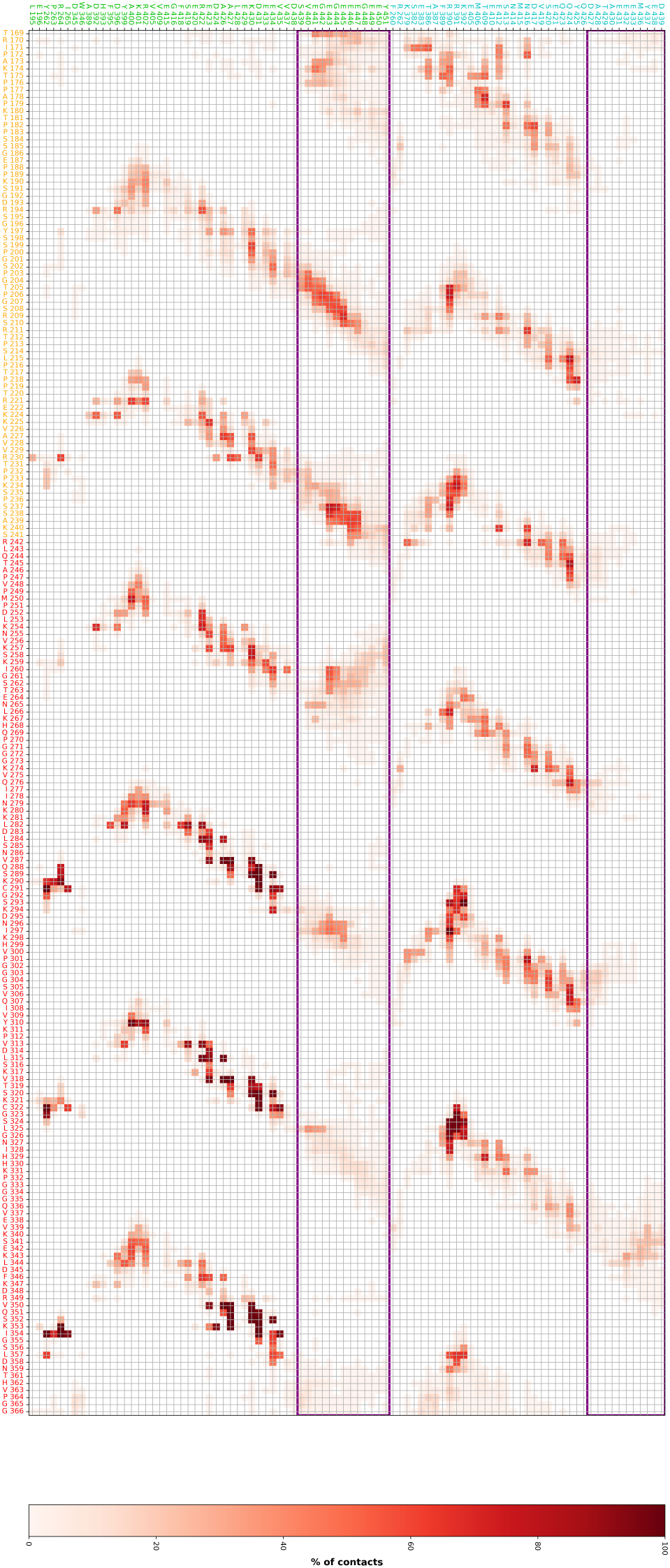
Contact map between the residues of the Tau fragment and tubulins *α*I/*β*III. Only residues making contact for at least 10% of the time are shown. *α*I-tubulin is in green, *β*III-tubulin in blue. Contacts with CTTs are boxed in purple. The PRR is in yellow and the repeat domains in red. The contact map of the *α*I/*β*I complex is available in Figure SI-6.

**Figure 9:**
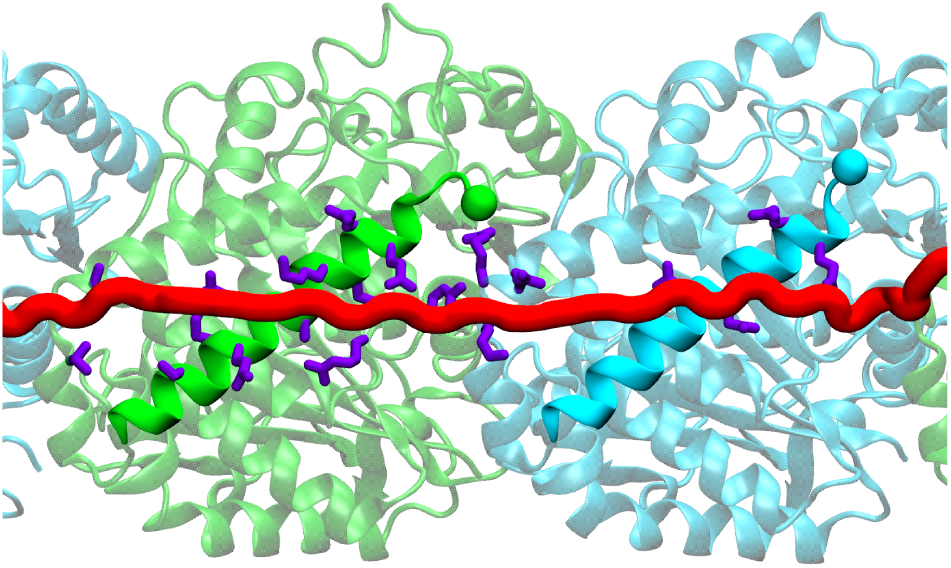
Upper view of the residues making contact with a residue of Tau at least 90% of the time. The H12 *α*-helix of tubulins and the Tau fragment are represented in opaque NewCartoon. CTTs were truncated for visibility and their base residue is represented by an opaque sphere.

Our complex extends further into the PRR region of Tau compared to the simulations of Brotzakis et al. since we added most of the P1 domain. This allowed us to notice that most of the P2 domain interacts more transiently with the tubulin cores than the repeats do (contacts never exceed 60% of the time). On the other hand, the P2 domain and in particular Tau residues Pro203-Ser210 and Ser237-Lys240 form contacts with the *α*-CTTs of monomers C and E respectively for a much longer time than any residue from the repeat domains. **It therefore seems that the stability of the Tau/PF complex is mediated by strong Tau/tubulin cores contacts for the repeat domains and by transient wrapping of the** *α***-CTTs for the P2 Tau domain.**

These observations are in agreement with previous experimental results which managed to obtain cryo-EM structures of repeat R2 and part of repeat R1 in complex with the tubulin cores.^43^ While the tubulin binding capacities of the PRR have been recognized,^12^ its interactions with tubulins have been characterized as fuzzier than those of the repeats.^13^ Further agreement is reached regarding the contact pattern with *α*-tubulin reported in Figure 8. Residues 224-KKVAVVR-230 form the majority of the contacts with the *α*-tubulin core, which was first reported for full Tau constructs in 1997 by Goode et al.^44^ and later confirmed for PRR peptides by Acosta et al.^45^ We can however assess from the contact maps that the contacts of the PRR with tubulins, while significant, are still transient (around 60% of simulated time). The involvement of the *α*-CTTs in the binding interactions of the PRR would thus reconcile all experimental observations, since the *α*-tubulin/PRR interface remains both significant yet fuzzy thanks to these CTTs. The important part played by the tubulin CTTs in Tau binding at the PRR level was also recently highlighted by Naveh-Tassa and Levy in their work modeling Tau-tubulin assemblies on the coarse-grained level. ^46^

#### b. Dynamic profile of Tau

A standard way to characterize the dynamics of an interface is to study its fluctuations compared to its native structure. We thus computed a modified RMSF of the Tau fragment where the sampled conformations of the entire complexes were first aligned based on the tubulin cores, then the fluctuations were calculated on the Tau monomer without any further alignment (Figure 10). This modified RMSF provides a sense of both the conformational heterogeneity of Tau, but also of the possible drift or detachments of the interface. For the repeat domains R3 and R4, the Tau fragments bound to the *α*I/*β*I and *α*I/*β*III PFs behave similarly, with a high mobility for the weak binding patches and a more stable interface for the strong binding patches. The C-ter part of the Tau fragment is highly mobile, with the last residues reaching values above 16 Å, probably due to a *truncation effect* as we do not simulate the entire length of Tau. For repeats R1 and R2, the two complexes behave differently, as the Tau monomer bound to the *α*I/*β*I PF experiences significant mobility at the strong binding patches of R1 and R2, while for the *α*I/*β*III complex the interface remains stable. The snapshots of Figure 10 illustrate how the SKIGS motif of R1 did not become buried at the interface between tubulins like the motif of R2 did, and how residues 263/264 interact with the base of the *α*I-CTT instead. This behavior is confirmed by the contact map (Figure SI-6), and is not observed for the *α*I/*β*III system. This instability of the R1 repeat is reminiscent of the difference observed in coarse-grained simulations of Tau, where we assessed that the local dynamics of the R1 repeat at its N-terminus was different from the other repeats.^47^ Overall, the PRR interface is oddly stable, while, according to NMR experiments by El Mammeri et al.^13^ and what was sampled by Brotzakis et al.,^16^ it should be very fuzzy. This could however be explained by the wrapping of the CTTs, and the RMSF is indeed slightly higher for the PRR than for the motifs.

**Figure 10:**
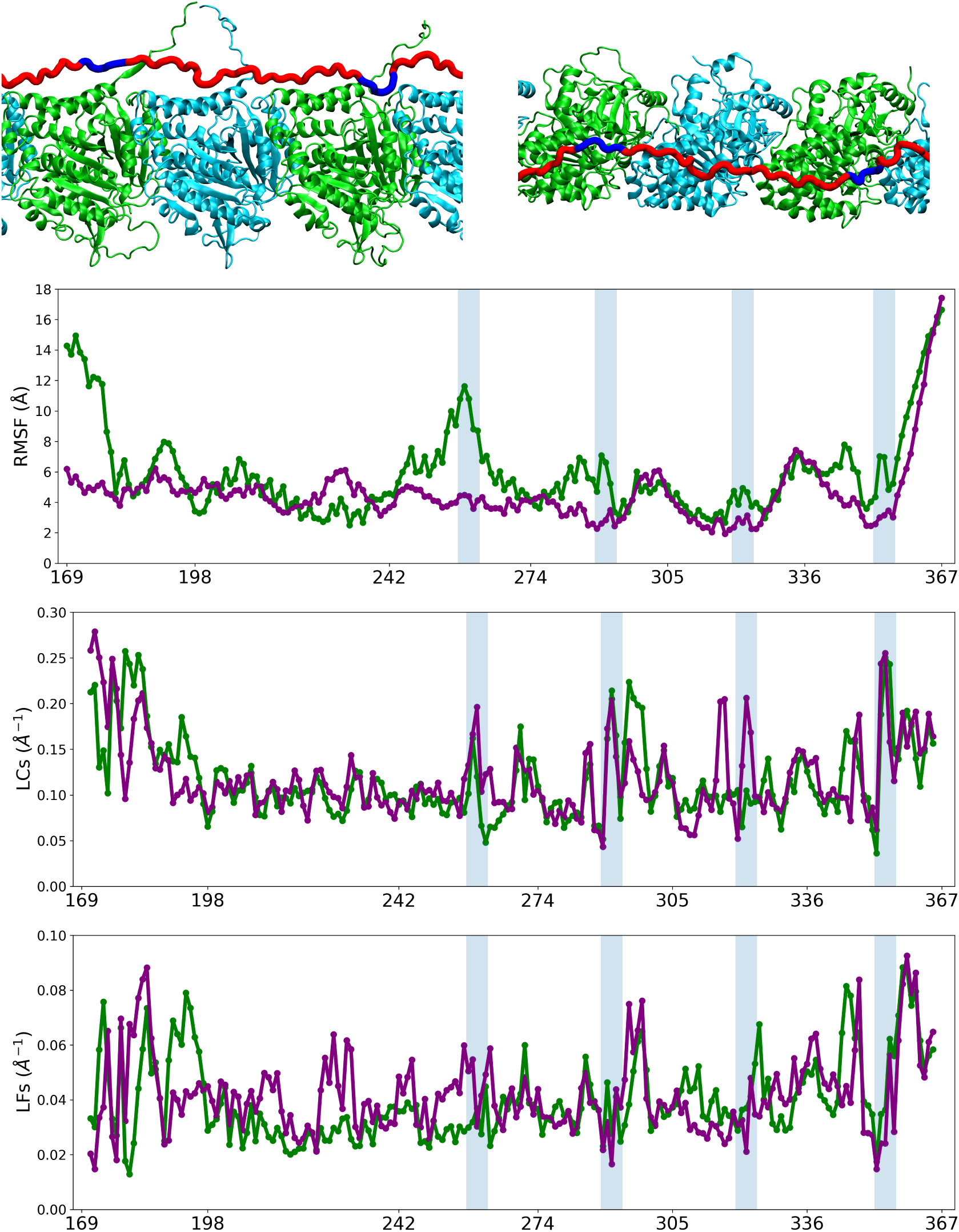
Snapshots of the side view (upper left) and top view (upper right) of the *α*I/*β*I system at 220ns, with the strong binding regions of R1 and R2 (residues 256-262 and 289-293) in dark blue. Upper row: RMSF of the Tau fragment with regards to the structure in the first frame. Middle row: Local Curvatures (LCs) of Tau. Bottom row: Local Flexibilities (LFs) of Tau. *α*I/*β*I is in green, *α*I/*β*III in purple. Shaded areas correspond to motifs SK[I/C]GS for the repeats.

The N-terminus for the Tau monomer bound to the *α*I/*β*I PF has a high RMSF similar to the C-terminus, indicating a large disruption with regards to the initial interface. We also noticed a non-negligible interaction between the N-terminal of the Tau fragment and the *α*I-CTT of monomer A for the *α*I/*β*III complex (Figure 8). These contacts do not resemble those made by other *α*-CTTs and are thus very likely arising from another *truncation effect* due to the section of Tau at Thr169. It is however reassuring for the dynamics of the AT8 epitope that the deviation in RMSF and the contacts with the *α*-CTT of monomer A occur from residues Thr169 to Glu187, seven residues away from the beginning of the epitope.

The differences between the *α*I/*β*I and the *α*I/*β*III complexes suggest that the interface is more stable when *β*III-CTTs are present. Considering that the *β*III isotype is more expressed in neurons where the Tau protein is also located,^48,49^ the stabilization of the Tau/PF interface suggested by the differences in RMSF with the *α*I/*β*I PF can be hypothesized to be a feature of the *β*III isotype.

While the modified RMSF informs on the deviations with respect to the original interface, LCs and LFs provide statistical information on the dynamics of the protein backbone at the residue level. For the repeat domains, the average curvature of the residues as described by LCs presents a peak for the central residue of each strong binding patch (Figure 10, middle row). LFs on the other hand mark a steep decrease for the motifs of R2, R3 and R4, coherent with the stabilization by burying in the intertubulin interface (Figure 10, bottom row). For the P2 domain, residues display stable values of LCs across the sequence between 0.07 and 0.13 Å*^−^*^1^, indicating that no burial at the intertubulin interface takes place for this domain. Surprisingly however, flexibility peaks can be observed for the *α*I/*β*I Tau monomer at residues 209 to 214 and 220 to 230. This attests to the fuzzy nature of the interface, as rigid backbone structures such as *α*-helices do not experience LF values higher than 0.02 Å*^−^*^1.32^

#### c. Comparison to other simulations highlight the *truncation effect*

##### Trimeric toy complex

In order to assess the effect of truncating the complex, we compared the contact maps obtained for the complexes containing only the R2 domain of Tau without the PGGG motif obtained in previous studies^18,19^ and those from the more complete Tau/PF complexes studied here (Figure SI-7). We expected the truncation to impact the N- and C-termini of the repeat, and this is indeed what we observe. However, even the central residues of the R2 domain were impacted by the truncation of the toy complexes, with variations in contact time with tubulins above 50% in gain or in loss. Notably, the *wrapping* phenomenon observed with the *α*I-CTT on the large complexes leads to wide discrepancies in the contact profiles, since in the toy complexes this CTT is unable to wrap due to the shortness of the Tau fragment. This indicates that truncating fuzzy complexes can have a profound impact on the interfaces involving IDPs, and should be considered in the interpretation of their dynamics.

##### 7PQC simulation by Brotzakis et al

We also computed the modified RMSF, LCs and LFs of the Brotzakis et al. complex based on the released trajectories (Figure SI-8). The RMSF shows slight peaks at mid-repeats, related to a large increase in LCs and LFs. The RMSF increases drastically at both termini, similarly to how it increases at the pseudo-repeat for our simulations. This suggests that this behavior might be a signature for a *truncation* effect, since it was observed for the R2/tubulins toy complex as well. ^18,19^ Furthermore, the RMSF is higher for residues 210 to 242 than in our complex. This can be explained by the quasi-absence of *wrapping* of the *α*-CTTs in the retrieved simulations of Brotzakis et al., which thus do not maintain the P2 domain close to the tubulin cores (see contact map on Figure SI-9). One could envision several reasons for this difference between our simulations. First, our tubulin sequences are not exactly similar, however the sequence of our *α*-CTTs is the same. Second, we employ different forcefields, but the similarity in CTT dynamics between our work and that of Laurin et al.^34^ suggests that this probably would not influence the conformational sampling, as discussed above.

We thus postulate that this absence of *wrapping* results from the difference in simulation techniques between our studies. Brotzakis et al. employed the EMMI method,^50^ which ultimately produces short concatenated replicas (32 in this case for an aggregate time of 400 ns). These would not leave enough time for the CTTs to rearrange and wrap themselves around Tau. These comparisons highlight the difficulties in probing large fuzzy systems. Employing incremental approaches in term of system size and complexity however did bring useful information on the complex, one should simply keep in mind the limitations induced by the *truncation* effect.

#### d. Tau influences the extension of the PF

In 2019, simultaneous studies by Siahaan et al. and Tan et al. revealed that Tau monomers could form a dense phase around microtubules termed at the time “condensates” or “islands”.^51,52^ Later in 2022, a follow-up study by Siahaan et al. further showed that this dense Tau phase effectively enveloped the microtubule, earning it the title of “Tau envelopes”.^9^ These Tau envelopes were shown to regulate the interaction with other MAPs such as kinesin with the microtubule, and to even cause a contraction of the microtubule lattice within them. A recent work by Beaudet et al. assessed the existence of Tau envelopes for in vivo neurons, and that phosphomimetic Tau mutants did not form such envelopes. ^10^ We thus designed the restraints on the tubulin cores so that the PF could expand and contract along its longitudinal axis (axis X on Figure 1). This allowed us to study its compaction with and without the Tau fragment, an effect that could not be investigated in the study of Brotzakis et al. because of the restraints imposed by the EMMI method on the tubulin cores.

We calculated the distance between the centers of mass of tubulin cores C and L as a function of time (Figure 11). An increase of more than 5 Å is observed during the first 100 ns for all simulations. However, this increase is slower and does not happen as sharply for the PFs with a Tau monomer bound to the tubulin cores than for those without. For the remaining 900 ns of the simulations, the distance between the COMs of the free tubulins remains higher than those bound by Tau, which is clearly visible when considering the distributions for each system (Figure 11B). The distribution of the distance for the *α*I/*β*I PF with Tau is lowered by 2 Å and its spread is diminished compared to the distribution with Tau. The difference is even more pronounced for the *α*I/*β*III PF, as the distribution is shifted by 4 Å. This represents a compaction with respect to the Tau-less complexes of around 1% of the lattice size. It should however be noted that we only model part of a single Tau monomer, which means that we probably miss on some of the cooperative effect induced by the high density of Tau monomers in an envelope, and a single PF rather than a full microtubule. Siahaan et al. reported that Tau monomers truncated between P2 and R1 did not form envelopes no matter the concentration in Tau protein.^51^ This indicates that the Tau-Tau interactions are mostly mediated by the projection region and the PRR, as suggested by Gamblin et al.^53^ Siahaan et al. also reported a compaction of 3.2 ± 1.1% over 59 envelopes, however the detail of each compaction event (Figure 1h of Ref ^9^) shows that smaller compactions, even of the order of 1%, are also observed. Our results therefore corroborate the role of Tau as a microtubule lattice regulator, and the compaction it induces in the PF could be part of its function as a microtubule stabilizer. Further investigations would require multiple Tau monomers to be added to the system, which is currently computationally untractable with all-atom simulations.

**Figure 11:**
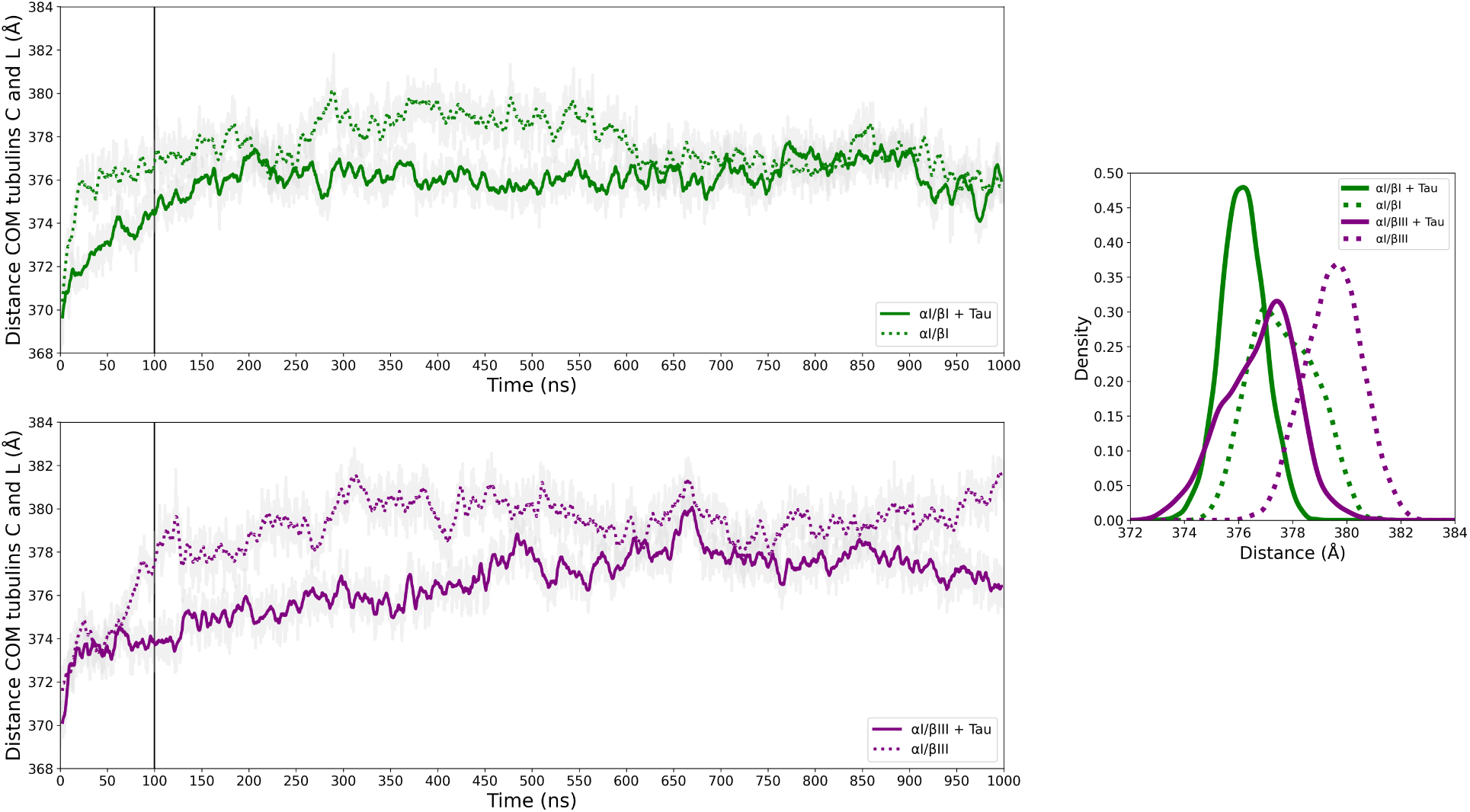
Left panels: Distance between the centers of mass of tubulin cores C and L as a function of time with and without Tau for the *α*I/*β*I PF (upper left) and the *α*I/*β*III PF (lower left). The black vertical line indicates the discarded 100ns that were excluded from the distributions on the right. Right panel: Density distribution of the distance between the centers of mass of tubulin cores C and L for all simulated systems.

## Conclusion

In this work, we provide an extensive characterization of the conformational ensembles of free CTTs from a tubulin protofilament. The periodicity of our PF reconstruction allowed for the sampling of 5 cumulative µs for each CTT type (*α*I with *β*I, *α*I with *β*III, *β*I and *β*III). We reveal a hook-like behavior of the tip of the *β*I and *β*III CTTs, loose for the former and tight for the latter. Upon Tau complexation, CTTs display different dynamic features and interactions depending on the region of Tau they are bound to. The consideration of their wrapping on the Tau fragment allowed us to define a region-dependent characterization of the Tau/tubulin interface: the repeat domains strongly interact with tubulin cores, burying their SKXGS motifs at the *α*/*β* interface, while the PRR region interacts more transiently with the cores but is stabilized by the wrapping of *α*I-CTTs. Tau impacts the mobility of the CTTs but also the PF itself, as we provided computational evidence of a contraction of the PF lattice upon Tau binding. This study is therefore an important contribution to our understanding of the Tau/tubulins interface and the disordered interplay which underlies their interactions. Furter work will investigate how multiple phosphorylations of Tau, which are connected to several neurodegenerative diseases, can impact both the Tau/PF interface and the mobility of the tubulin CTTs.

## Acknowledgement

This work was supported by the ANR (MAGNETAU-ANR-21-CE29-0024) and the “Initiative d’Excellence” program from the French State (Grants “DYNAMO”, ANR-11-LABX-0011, and “CACSICE”, ANR-11-EQPX-0008). This project was provided with computing HPC and storage resources by GENCI (project A0170711021).

## Supporting Information Available

The following files are available free of charge.

- Additional data regarding the simulated systems composition, the CTTs DSSPs, Tau/tubulin contact maps, CTTs curvature and mobility are available as supplementary information.
- All MD trajectories without the solvent and their topologies are deposited in Zenodo: https://zenodo.org/records/19886983

