## Supplementary information for "Simulations of an extended Tau/tubulins interface reveal a complex disorder-disorder interplay mediated by the C-terminal tails"

Table S1: Composition of the simulated systems. Bulk ions are chloride and potassium. Ligands are GDP, GTP and their associated magnesium cations.

| | **Total** | **Protein** | **Water molecules** | **Bulk ions** | **Ligand atoms** |
|---|---|---|---|---|---|
| **$\alpha$I/$\beta$I PF + Tau** | 1237769 | 99308 | 378475 | 2486 | 550 |
| **$\alpha$I/$\beta$I PF** | 1237911 | 96327 | 379506 | 2516 | 550 |
| **$\alpha$I/$\beta$III PF + Tau** | 1238027 | 99993 | 378333 | 2485 | 550 |
| **$\alpha$I/$\beta$III PF** | 1238263 | 97012 | 379396 | 2513 | 550 |

Table S2: Frames (out of 10000) of the simulation for the $\alpha$I CTTs in the $\alpha$I/$\beta$I complex during which a transversal, longitudinal, or both *wrapping* of the CTT around Tau is present

| | **C** | **E** | **G** | **I** | **K** |
|---|---|---|---|---|---|
| **transversal** | 160:600 | 20:2770<br>8120:9320 | 5030:5240 | 380:620<br>5850:6210 | 0 |
| **longitudinal** | 0 | 7480:7670<br>9770:10000 | 0 | 0 | 1350:1670<br>4980:5380<br>8190:10000 |
| **both** | 1240:10000 | 2820:5950<br>6250:6640<br>7130:7350 | 2320:2840 | 0 | 0 |

Table S3: Frames (out of 10000) of the simulation for the $\alpha$I CTTs in the $\alpha$I/$\beta$III complex during which a transversal, longitudinal, or both *wrapping* of the CTT around Tau is present

| | **C** | **E** | **G** | **I** | **K** |
|---|---|---|---|---|---|
| **transversal** | 120:1320<br>1840:2780 | 0 | 9800:9950 | 0 | 7180:7450 |
| **longitudinal** | 5120:5350 | 640:980<br>3290:5450<br>5680:10000 | 500:1320<br>4540:7170 | 0 | 0 |
| **both** | 5390:10000 | 1430:3170 | 0 | 4100:6150 | 1700:2780 |

Table S4: Frames (out of 10000) of the simulation for the $\beta$I CTTs in the $\alpha$I/$\beta$I complex during which a transversal, longitudinal, or both *wrapping* of the CTT around Tau is present

| | **D** | **F** | **H** | **J** | **L** |
|---|---|---|---|---|---|
| **transversal** | 0 | 3480:4330 | 7890:8570 | 0 | 0 |
| **longitudinal** | 0 | 0 | 0 | 0 | 0 |
| **both** | 0 | 0 | 0 | 0 | 0 |

Table S5: Frames (out of 10000) of the simulation for the $\beta$III CTTs in the $\alpha$I/$\beta$III complex during which a transversal, longitudinal, or both *wrapping* of the CTT around Tau is present

| | **D** | **F** | **H** | **J** | **L** |
|---|---|---|---|---|---|
| **transversal** | 0 | 0 | 0 | 0 | 7130:7310<br>8650:9100 |
| **longitudinal** | 0 | 0 | 0 | 0 | 0 |
| **both** | 0 | 0 | 0 | 0 | 0 |

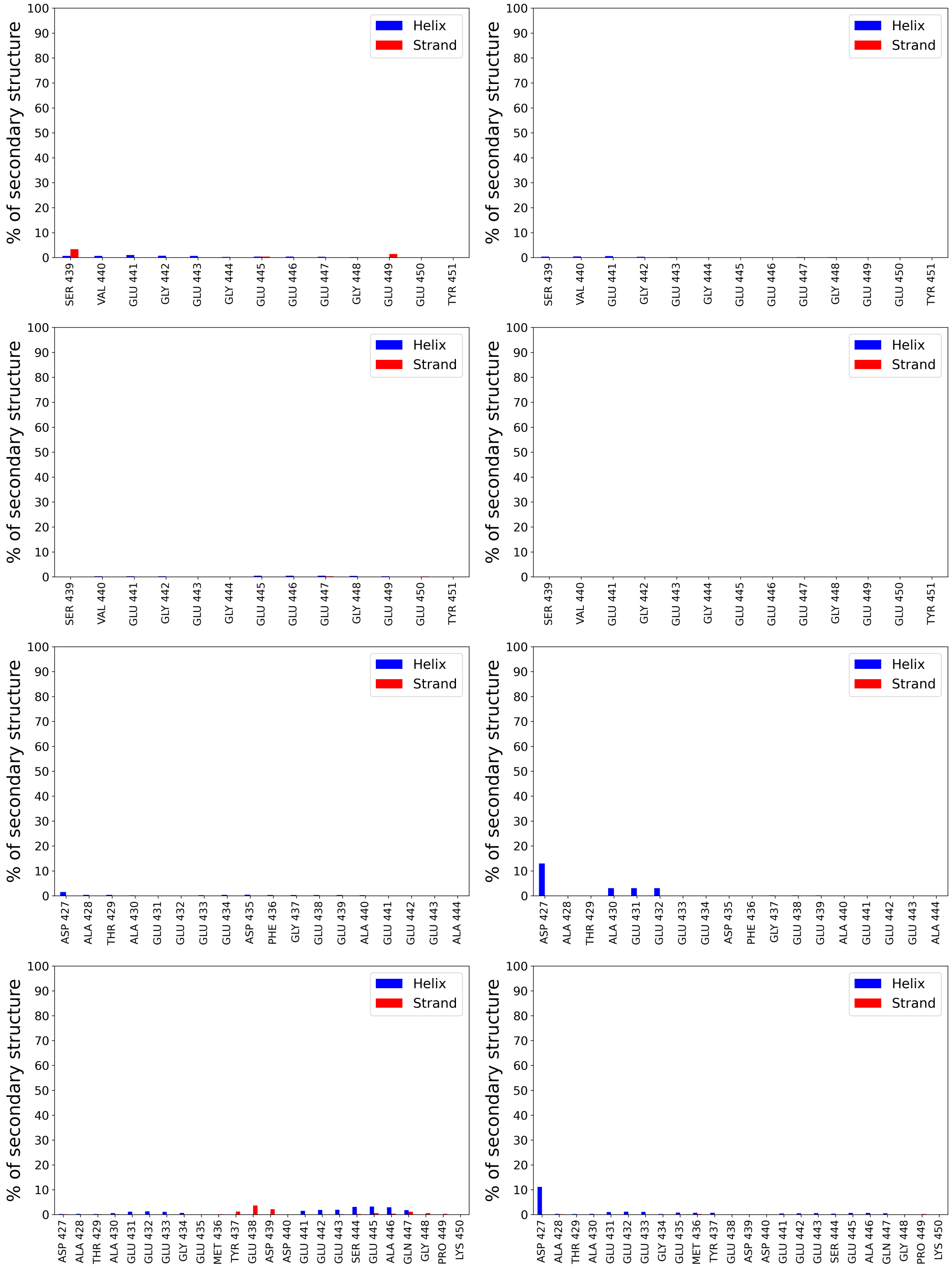


Figure S1: Secondary structure propensities of the CTTs with (right column) and without (left column) Tau. First row: $\alpha$I CTTs in the $\alpha$I/$\beta$I complex. Second row: $\alpha$I CTTs in the $\alpha$I/$\beta$III complex. Third row: $\beta$I CTTs. Fourth row: $\beta$III CTTs

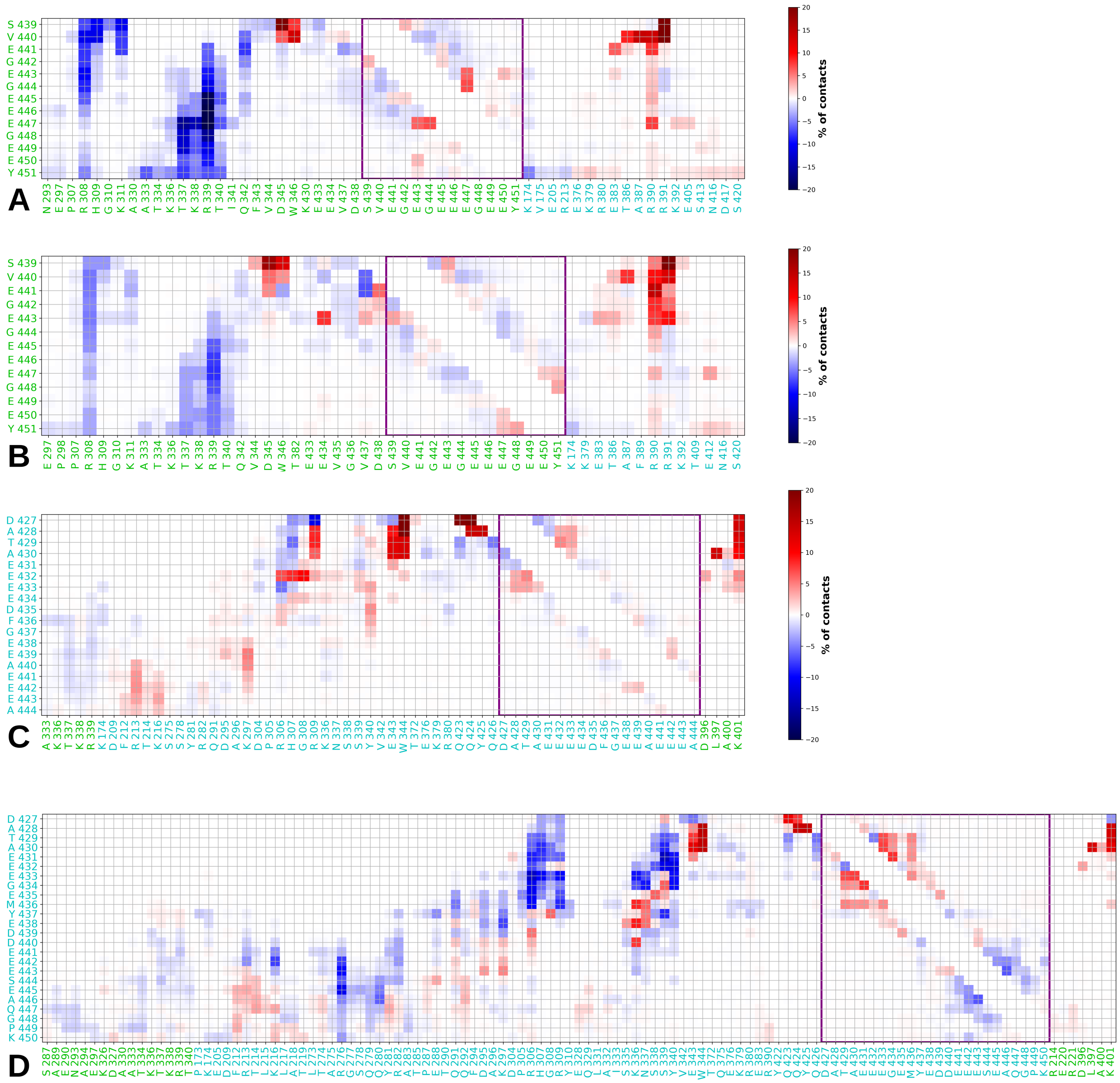


Figure S2: Concatenated difference of contacts between the CTTs and tubulin $\alpha$ (in green) and $\beta$ (in blue) with and without Tau. Tubulin residues that did not contact the CTT at least once during the simulations were filtered out. A) $\alpha$I CTTs in the $\alpha$I/$\beta$I complex. B) $\alpha$I CTTs in the $\alpha$I/$\beta$III complex. C) $\beta$I CTTs. D) $\beta$III CTTs

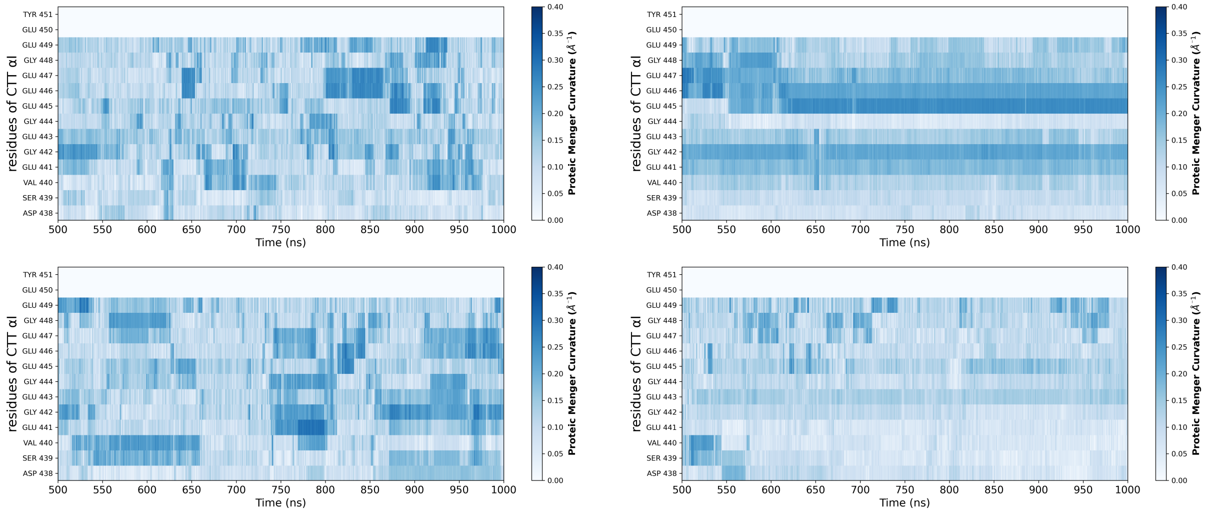


Figure S3: Proteic Menger Curvatures (PMCs) as a function of simulation time of the $\alpha$I-CTT of monomer C. Upper row: CTT in the $\alpha$I/$\beta$I PF. Lower row: B) CTT in the $\alpha$I/$\beta$III PF. Left column is the CTT in the Tau-less complex, right column is the CTT in contact with Tau.

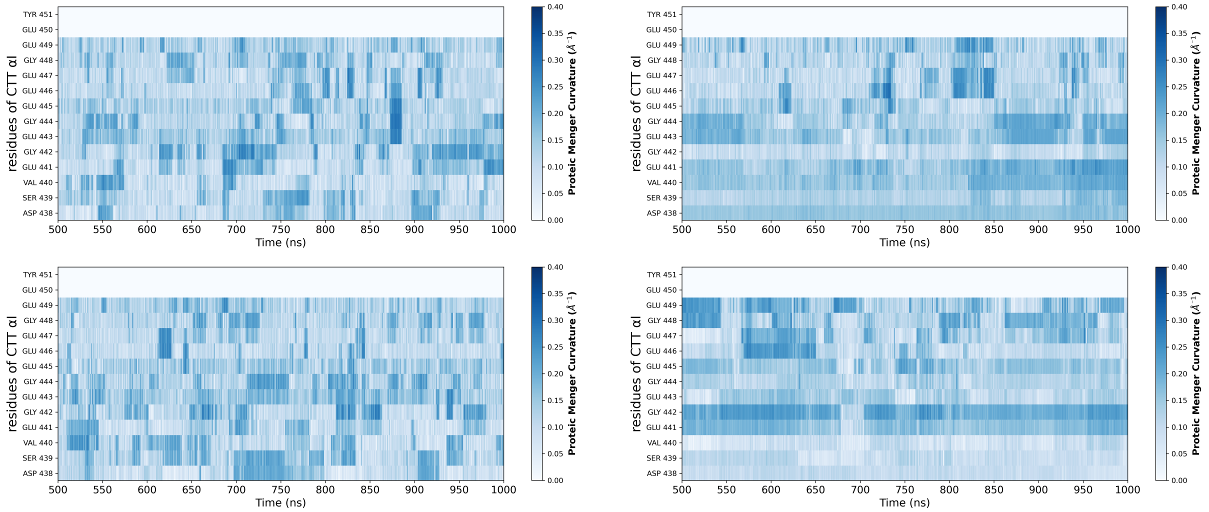


Figure S4: Proteic Menger Curvatures (PMCs) as a function of simulation time of the $\alpha$I-CTT of monomer E. Upper row: CTT in the $\alpha$I/$\beta$I PF. Lower row: B) CTT in the $\alpha$I/$\beta$III PF. Left column is the CTT in the Tau-less complex, right column is the CTT in contact with Tau.

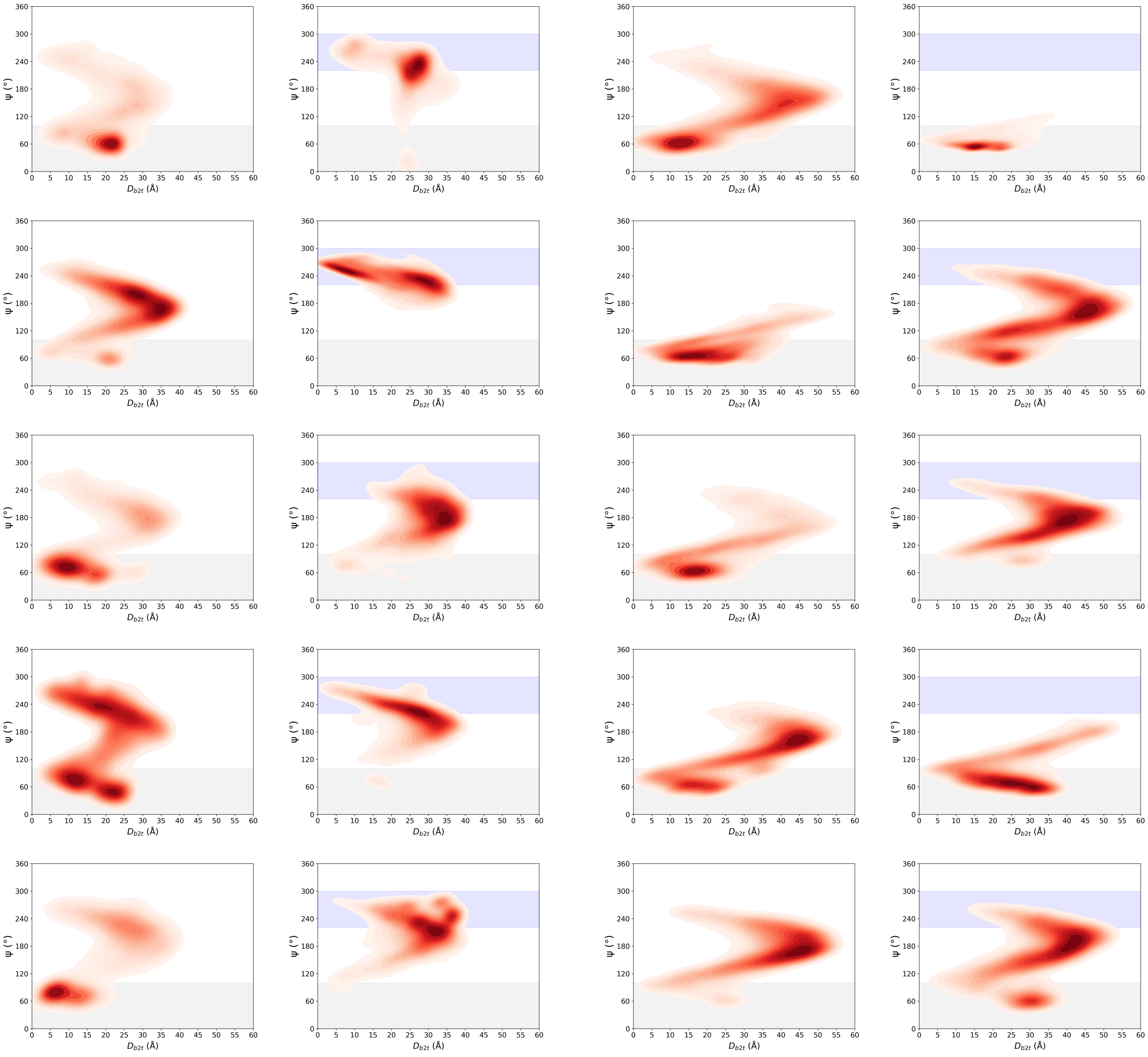


(a) $\alpha$I-CTTs

(b) $\beta$III-CTTs

Figure S5: Landscapes of the $\Psi_{CTT}/D_{b2t}$ values for the CTTs of the bare $\alpha$I/$\beta$I PF (left columns of S5a and S5b) and the PF in complex with the Tau fragment (right columns of S5a and S5b). From top to bottom, plots correspond to monomers C, E, G, I and K for the $\alpha$I-CTTs and monomers D, F, H, J and L for the $\beta$III-CTTs. The range of $\Psi_{Tau}$ values is shaded in blue. The range of $\Psi_{CTT}$ values corresponding to a position that would be buried into the inter-PF interface is shaded in grey.

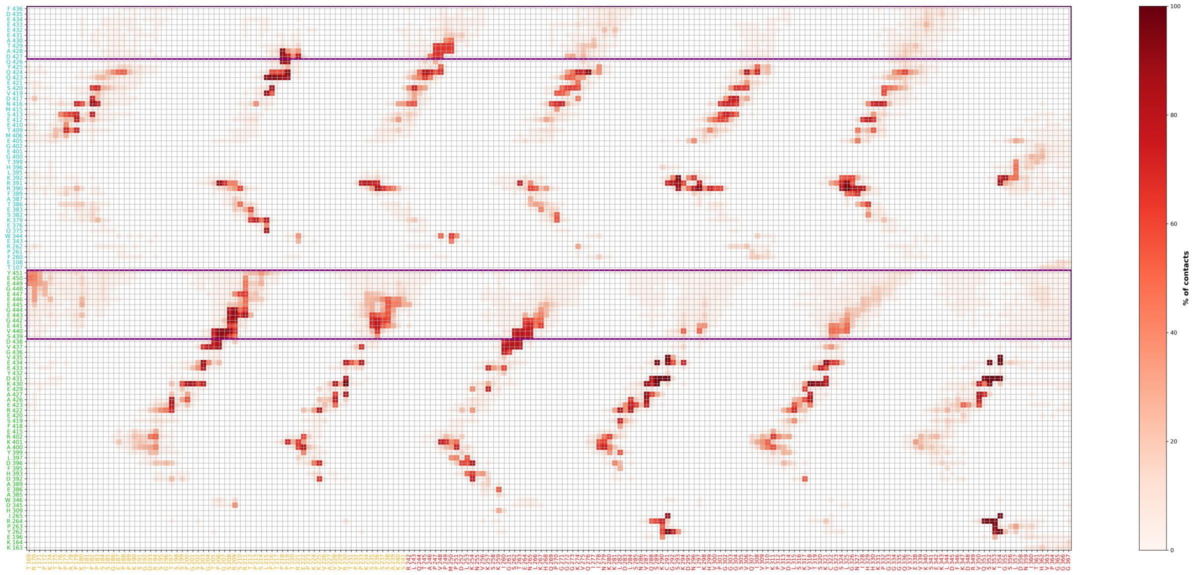

Figure S6: Contact map between the Tau fragment and tubulins $\alpha$I/$\beta$I. Only residues making a contact for at least 10% of the time are shown. $\alpha$I-tubulin is in green, $\beta$I-tubulin in blue. Contacts with CTTs are boxed in purple. The PRR is in yellow and the repeat domains in red.

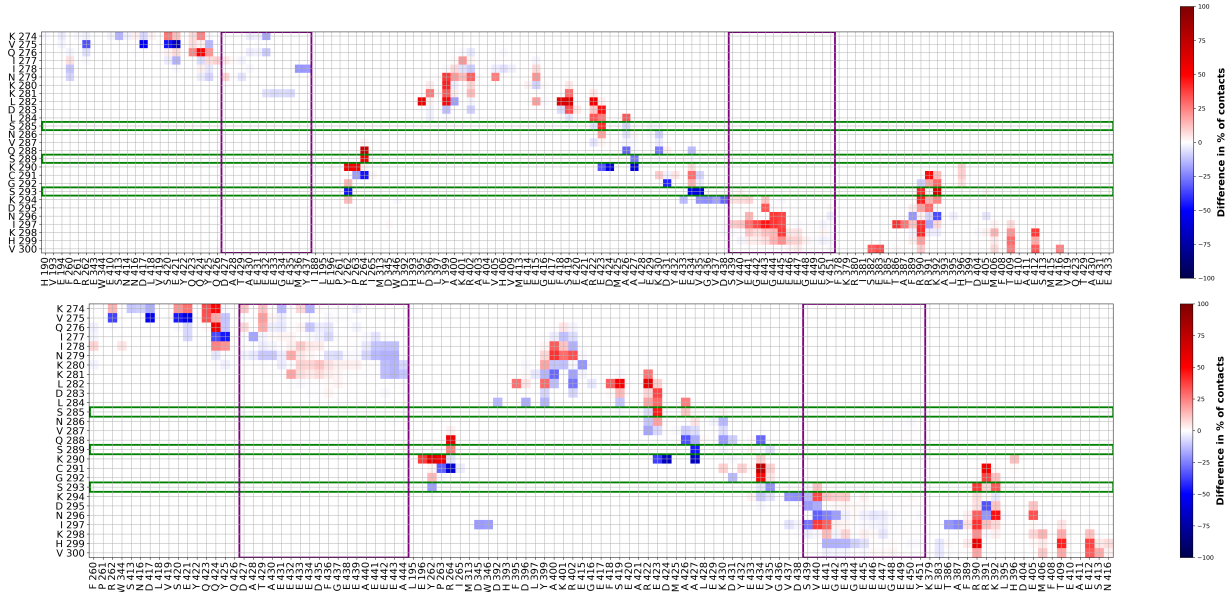

Figure S7: Difference of contacts Tau/tubulins between the full-size system and the trimer simulated in Ref,[1] taking the trimer as a reference. Tubulin residues that did not contact the CTT at least once during the simulations were filtered out. Upper panel: $\alpha$I/$\beta$I. Lower panel: $\alpha$I/$\beta$III.

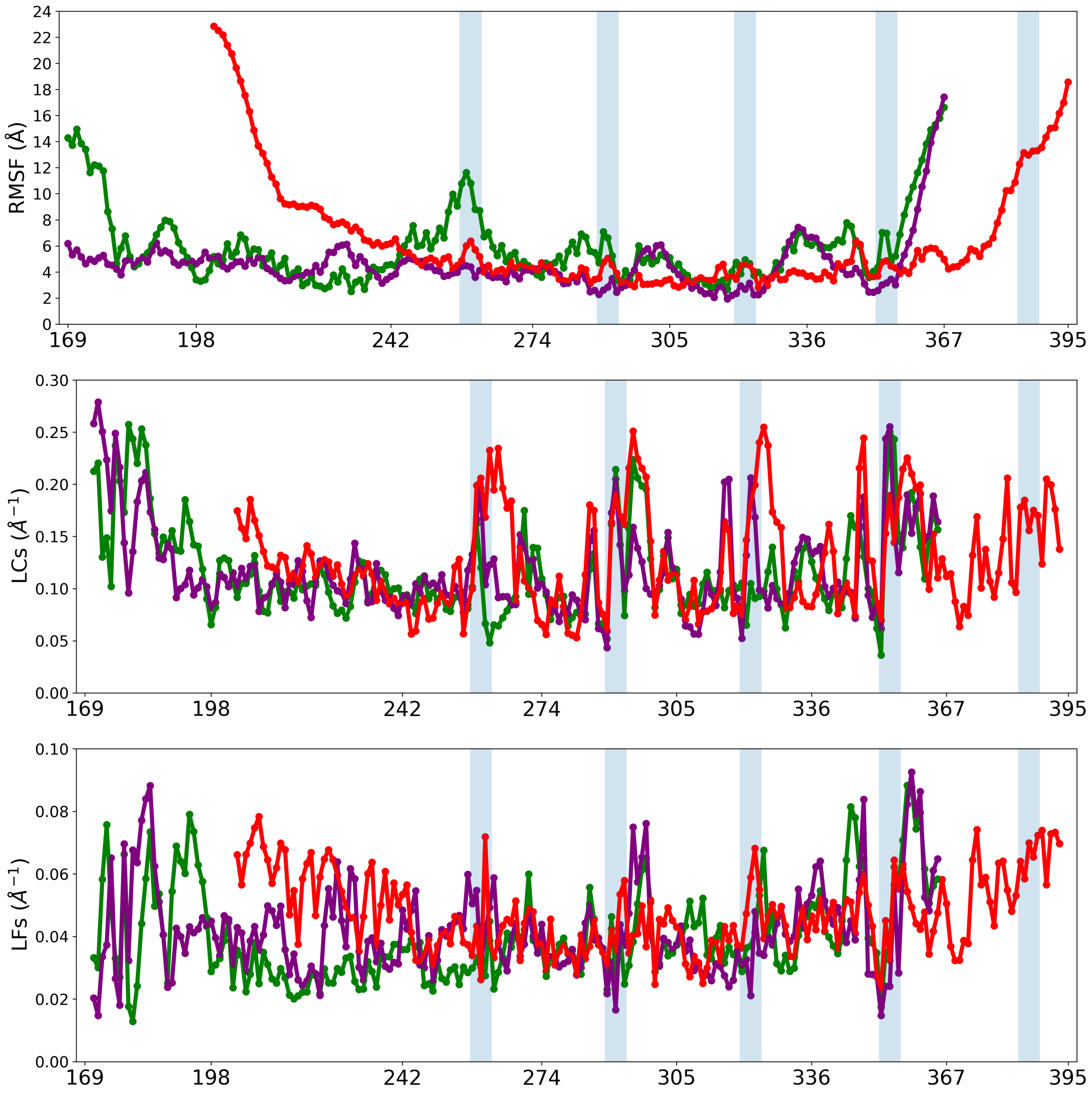


Figure S8: Comparison of the modified RMSF, LCs and LFs from Brotzakis et al. (in red) with our simulations. Upper row: RMSF of the Tau fragments with regards to the structure in the first frame. Middle row: Local Curvatures (LCs) of Tau. Bottom row: Local Flexibilities (LFs) of Tau. $\alpha$I/$\beta$I is in green, $\alpha$I/$\beta$III in purple. Shaded areas correspond to motifs SK[I/C]GS for the repeats.

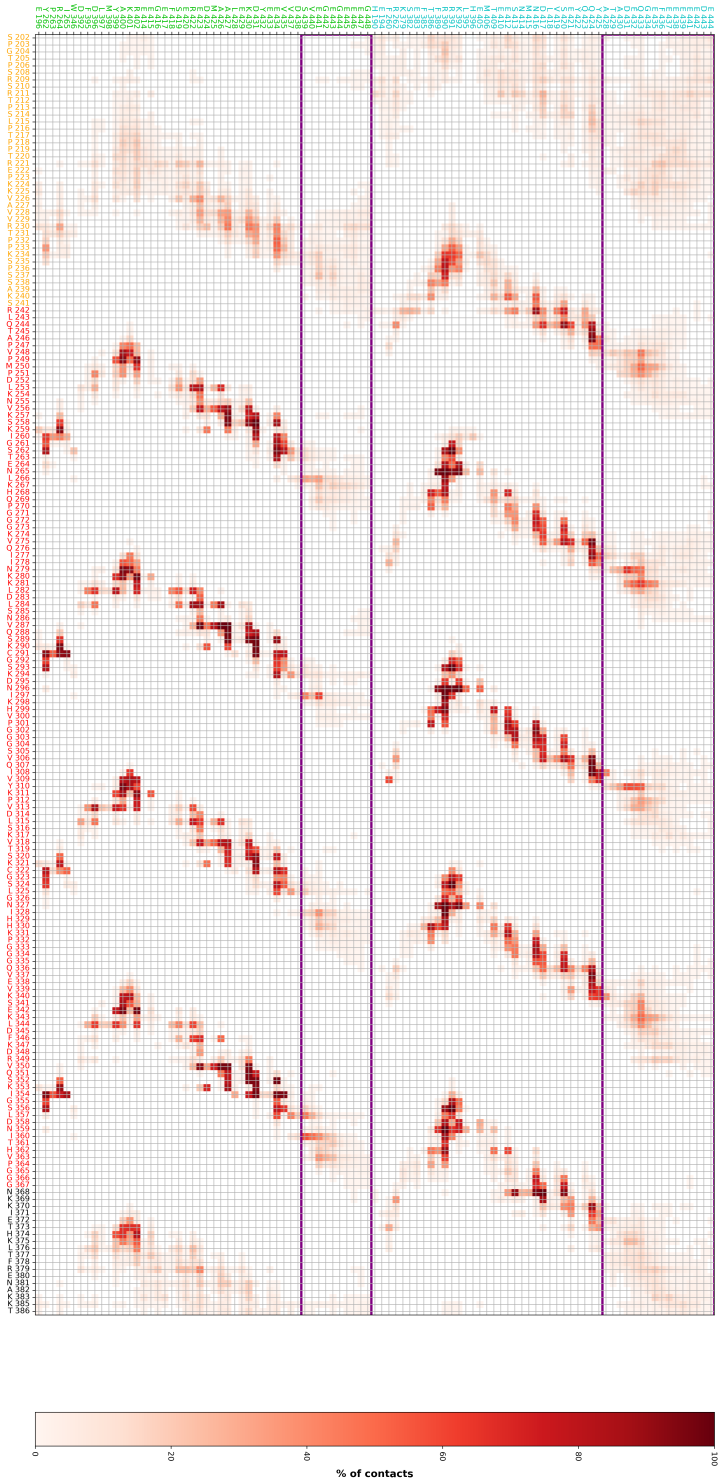

Figure S9: Contact map between the tubulins and the Tau fragment simulated by Brotzakis et al.[2] Only residues making a contact for at least 10% of the time are shown. $\alpha$I-tubulin is in green, $\beta$III-tubulin in blue. Contacts with CTTs are boxed in purple. The PRR is in yellow, the repeat domains R1/R2/R3/R4 in red and the pseudorepeat R' in black.
